# Moonlighting Role of Meiotic SYCP1 in Breast Cancer: A Chromatin-Bound Regulator of DNA Repair, Transcription, and Drug Resistance

**DOI:** 10.1101/2025.09.18.677087

**Authors:** LC Brennan, OV Grinchuk, M Pachon, IF Sou, CJ Fawcett, CG Nogueira, M Guthrie, AD Bates, M Hine, A Thomaz, Z Hu, AB Fielding, OR Davies, W-W Tee, UL McClurg

## Abstract

Maintenance of genome integrity is essential for cellular homeostasis, and its perturbation leads to tumorigenesis. Here, we uncover an unanticipated somatic role for the synaptonemal complex protein SYCP1—previously regarded as strictly meiosis-specific—in a broad spectrum of human cancers including breast cancer. Through integrative genomic, proteomic, and functional analyses, we demonstrate that SYCP1 is aberrantly re-expressed in tumor cells, where it actively promotes DNA damage repair, cell cycle progression, and malignant growth. SYCP1 binds chromatin at regulatory elements and directly controls transcriptional programs governing genome maintenance, including key effectors such as *CCNB1*, *PCNA*, *RAD51C*, and *H2AX*. Loss of SYCP1 impairs DNA repair kinetics, attenuates tumor cell proliferation and migration, and increases sensitivity to chemotherapeutics cisplatin and gemcitabine. Mechanistically, SYCP1 interfaces with chromatin remodeling complexes and transcription factors SP1 and SP2, modulating their genomic occupancy and facilitating oncogenic transcriptional outputs. Clinically, high SYCP1 expression stratifies patients with poor prognosis and therapy resistance across multiple cancer types. Our findings illuminate a previously unrecognized moonlighting function of SYCP1 in somatic cancer cells and position it as a critical chromatin-associated regulator of genome stability, with implications for biomarker development and therapeutic targeting.

## Introduction

Preserving genome integrity is fundamental to the homeostasis and survival of somatic cells. Genomic instability is a defining hallmark of cancer and a major contributor to malignant transformation and therapeutic resistance [1]. Similarly, genome stability is indispensable during gametogenesis, where meiosis imposes highly orchestrated chromosomal movements to ensure accurate homolog pairing and recombination. Unlike mitosis, meiosis introduces programmed DNA double-strand breaks (DSBs) to facilitate homologous recombination and crossover formation—processes that are essential for genetic diversity and proper chromosome segregation. This added layer of complexity is stabilized by the synaptonemal complex (SC), a supramolecular protein scaffold that mediates synapsis and promotes crossover maturation [2]. Perturbations in SC assembly cause infertility [3].

In stark contrast, somatic cells must avoid recombination instability. The presence of haploid-like gene expression or unregulated DNA breakage in somatic tissues has deleterious consequences, causing chromosomal instability and accelerating tumorigenesis [4]. To preserve genomic integrity, meiotic genes—particularly those involved in chromosome structure and recombination—are stringently silenced in somatic lineages. The human SC comprises eight autosomally encoded proteins— SYCP1, SYCP2, SYCP3, SYCE1, SYCE2, SYCE3, TEX12, and SIX6OS1—all canonically restricted to germ cells [2]. However, recent studies reveal aberrant re-expression of SC components in diverse human cancers, classifying them among cancer-testis antigens (CTAs) or more broadly as germ cell cancer genes (GCCGs) [4–9]. Given their pivotal roles in meiotic chromosome dynamics, the reactivation of SC proteins in somatic cancer cells represents a profound threat to genome stability, potentially fueling malignant evolution.

Cancer cells frequently co-opt developmental programs to achieve proliferative plasticity and survive environmental stressors. The reactivation of meiotic genes exemplifies this strategy, allowing tumors to exploit non-somatic machineries for adaptive advantage. In this context, we identify SYCP1—the central transverse filament of the SC [10]—as a somatically reactivated protein with direct oncogenic potential. SYCP1 is repurposed in cancer cells to promote DNA repair, sustain proliferation, and drive resistance to genotoxic therapies.

Our data reveal that SYCP1 expression is a functional mediator of cancer cell fitness. In tumors, SYCP1 facilitates DNA damage repair and rewires transcriptional programs essential for cell cycle progression. Mechanistically, SYCP1 binds to gene promoters and chromatin regulatory regions, modulating the expression of critical DNA repair and replication factors, including *CCNB1*, *PCNA*, *RAD51C*, and *H2AX*. Loss of SYCP1 impairs these processes, abrogates cellular proliferation and migration, and sensitizes tumor cells to DNA-damaging agents such as gemcitabine and cisplatin.

Strikingly, SYCP1’s role in cancer diverges from its canonical function in meiosis. In the germline, SYCP1 acts as a structural scaffold aligning homologous chromosomes to support recombination fidelity [11]. In contrast, in the somatic context of cancer, SYCP1 operates as a chromatin-bound regulator of gene expression. We identify direct interactions between SYCP1 and chromatin remodeling factors and show that its loss disrupts the chromatin recruitment of transcriptional regulators SP1 and SP2. This suggests that SYCP1 is co-opted as a transcriptional effector, linking chromatin structure to oncogenic gene expression.

Because meiosis and mitosis operate in fundamentally distinct cellular environments— with divergent proteomes, epigenetic landscapes, and regulatory cues—meiotic proteins like SYCP1 may acquire novel functions when ectopically expressed in somatic cancer cells. Rather than merely reinstating meiotic processes, these proteins moonlight in oncogenic pathways, driving tumor progression through mechanisms distinct from their original biological context. Our findings uncover a previously unappreciated role for SYCP1 as a chromatin-associated modulator of genome stability in cancer, positioning it as both a mechanistic driver of tumor evolution and a candidate therapeutic target.

## Results

### SYCP1 is commonly reactivated in cancer patients

While prior studies have reported aberrant re-expression of *SYCP1* at the transcript level across various malignancies [12–20], the functional implications of this reactivation— and whether it results in detectable protein expression—have remained largely unresolved. To address this, we conducted a comprehensive analysis of *SYCP1* expression in patient-derived datasets and cancer cell lines.

Systematic interrogation of The Cancer Genome Atlas (TCGA) revealed that *SYCP1* mRNA is recurrently upregulated across a wide array of solid tumors, with particularly high expression observed in lung and breast cancers. Notably, approximately one-in-twenty of patients with these malignancies exhibited marked *SYCP1* mRNA overexpression (**Fig. 1a**). Importantly, elevated *SYCP1* expression significantly correlated with reduced overall survival in multiple cancer types, including cholangiocarcinoma, thymoma, skin cutaneous melanoma, uterine corpus endometrial carcinoma, and rectal and gastric adenocarcinomas (***Suppl. Fig. 1***). A similar negative association was observed with relapse-free survival (**Suppl. Fig. 2**), suggesting that *SYCP1* expression may serve as a clinically relevant prognostic biomarker.

**Figure 1.**
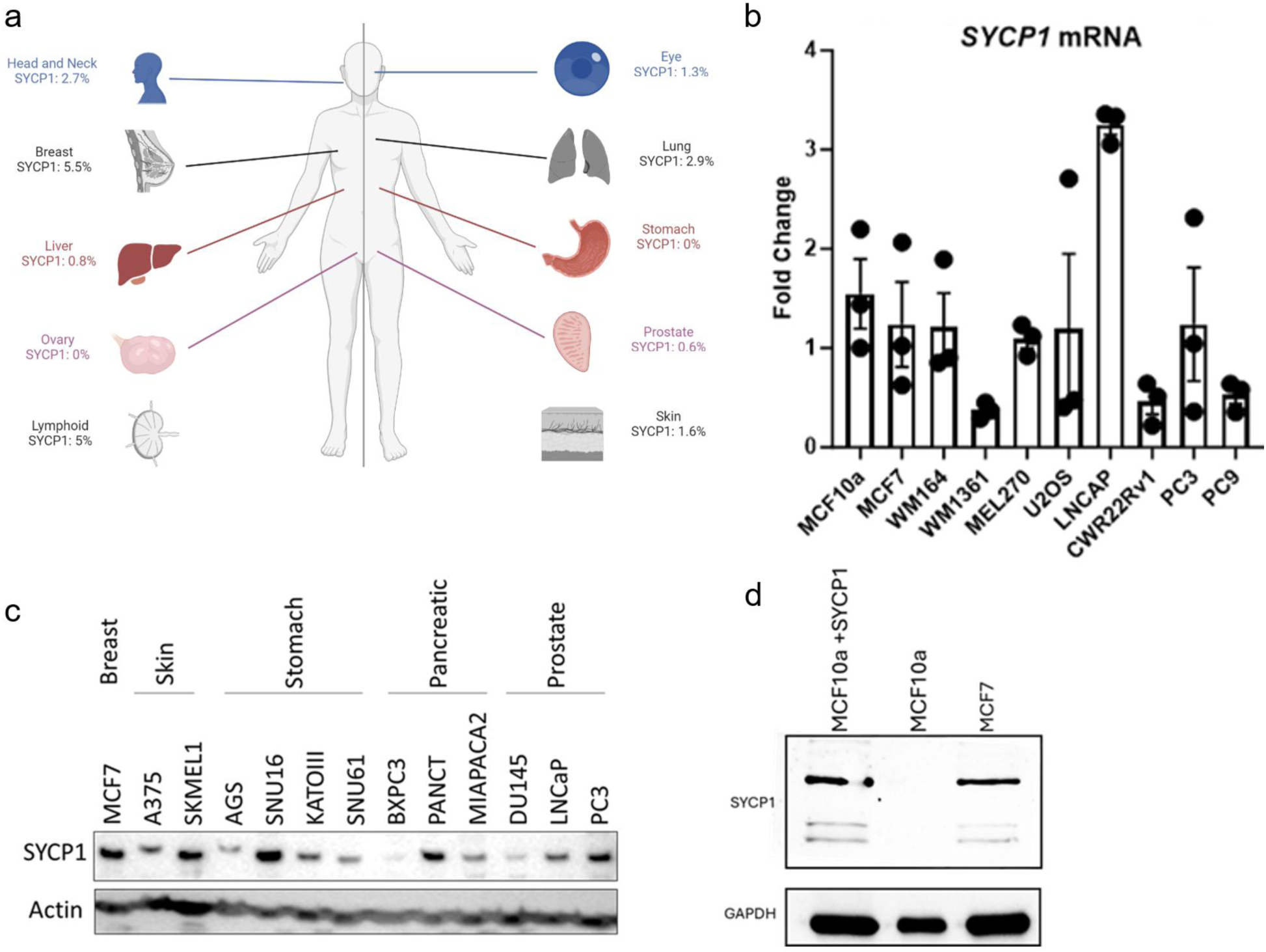
SYCP1 is commonly expressed in cancer. (**a**) TCGA cancer patient atlas was analysed to determine % of cancer patients with SYCP1 mRNA expression. (**b**) SYCP1 mRNA levels in cancer cell lines analysed by qPCR. MCF10a non transformed breast epithelium, MCF7 breast cancer, WM164 and WM1361 melanoma, Mel270 uveal melanoma, U2OS osteosarcoma, LNCAP, CWR22Rv1 and PC3 prostate cancer, PC9 lung cancer. (**c**) SYCP1 protein levels across cancer cell lines. (**d**) SYCP1 protein levels in breast epithelial cells including MCF10a cells following SYCP1 transfection (+SYCP1).

To experimentally validate these observations, we assessed *SYCP1* expression across a panel of commonly used cancer cell lines. Consistent with patient data, the majority of tested models displayed robust expression of *SYCP1* at both the transcript (**Fig. 1b**) and protein levels (**Fig. 1c**, **Suppl. Fig. 3**). Among these, MCF7 breast cancer cells exhibited particularly high levels of SYCP1 protein, while no detectable expression was observed in the non-transformed MCF10A breast epithelial line (**Fig. 1d**). These findings underscore the cancer-specific nature of *SYCP1* expression and highlight MCF7 cells as a relevant model for mechanistic interrogation.

Interestingly, our data also revealed a weak correlation between *SYCP1* mRNA and protein levels, suggesting significant post-transcriptional regulation. This observation emphasizes the importance of directly assessing protein expression when evaluating the reactivation of meiotic genes in cancer. Overall, these results establish that *SYCP1* is not only re-expressed in cancer at the mRNA level but also translated into protein in a subset of tumors, where it may play a functional role in tumor progression.

### SYCP1 binds to chromatin in cancer cells

To elucidate the functional role of aberrantly expressed SYCP1 in cancer, we first examined its subcellular localization. Immunofluorescence analysis revealed that SYCP1 localizes predominantly to the nucleus in MCF7 breast cancer cells (**Fig. 2a**) and in additional cancer cell models (**Suppl. Fig. 4a–c**). These findings were corroborated by biochemical cell fractionation, which confirmed nuclear enrichment of endogenous SYCP1 (**Fig. 2b**, **Suppl. Fig. 4d**).

**Figure 2.**
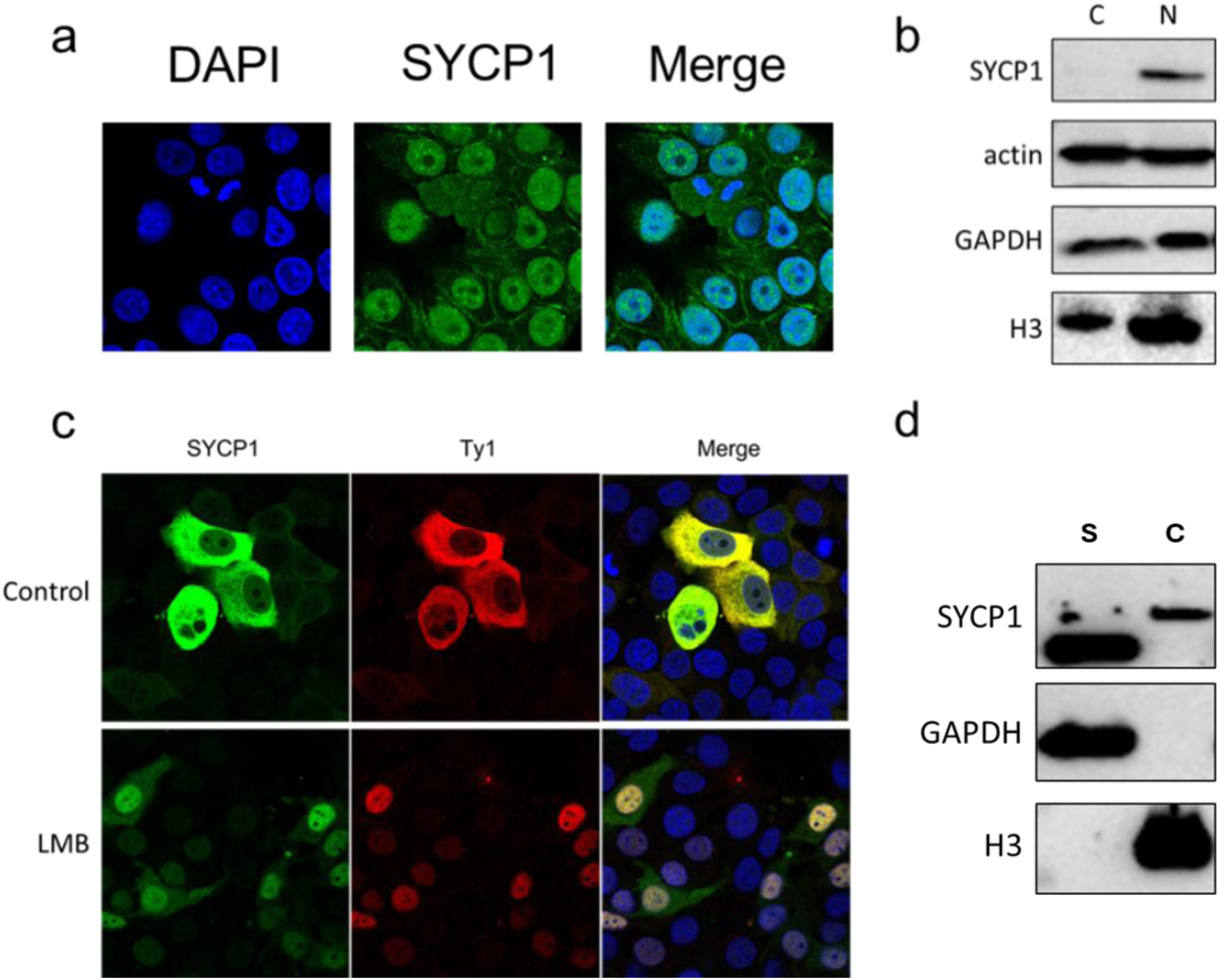
In cancer cells SYCP1 localises to the nucleus where it binds to the chromatin. (**a**) MCF7 cells with immuno-labelled SYCP1. (**b**) Cell fractionation of MCF7 cells to cytoplasm (C) and nucleus (N). (**c**) MCF7 cells transfected with dox inducible SYCP1-3xTy1 vector and SYCP1 overexpression was induced upon treatment with 5 μg/ml doxycycline. Where indicated cells were treated with Leptomycin B for 3 hours. (**d**) MCF7 breast cancer cells were fractionated to chromatin (C) and the soluble fraction containing cytoplasm and nucleoplasm (S).

To investigate the behavior of SYCP1 in a controlled system, we generated a Doxycycline (Dox)-inducible SYCP1 construct fused to a 3xTy-1 epitope tag and established a stable MCF7 cell line expressing the tagged protein. Upon induction, SYCP1 localized to both the cytoplasm and the nucleus, with a pronounced cytoplasmic predominance under steady-state conditions. However, pharmacological inhibition of nuclear export resulted in exclusive nuclear retention of SYCP1 (**Fig. 2c**), indicating that SYCP1 undergoes active nucleocytoplasmic shuttling in cancer cells.

To determine whether nuclear SYCP1 is associated with chromatin, we performed chromatin fractionation assays. These revealed that a substantial pool of endogenous SYCP1 is present in the chromatin-bound fraction, alongside a soluble nuclear fraction (**Fig. 2d**). Notably, the chromatin-associated form of SYCP1 exhibited a distinct electrophoretic mobility shift consistent with post-translational modification, suggesting that chromatin binding may be regulated by specific molecular cues.

Together, these findings demonstrate that SYCP1 is not only reactivated in cancer cells but is actively trafficked to the nucleus, where it engages with chromatin. This subcellular localization pattern supports a functional role for SYCP1 in regulating gene expression and chromatin dynamics in tumor cells.

### SYCP1 binds to the promoters of genes responsible for DNA damage response and cell cycle

To investigate the role of chromatin-bound SYCP1, we performed Cleavage Under Targets & Tagmentation (CUT&Tag) profiling [21] in the SYCP1-3xTy1 MCF7 Dox-inducible cell line, using a Ty1 targeting antibody to map SYCP1 genomic binding sites (**Fig 3**). A total of 8432 peaks were detected and we found that SYCP1 predominantly binds to gene promoters (approx. 68%, **Fig 3a and 3b**). Notably, SYCP1 was significantly enriched at promoters of genes responsible for transcription factor binding, RNA metabolism and processing, cell cycle, DNA repair and response to stimuli (**Fig 3c**). Of particular interest was SYCP1 localization to the promoters of key cell cycle genes (e.g., *CCNB1* and *PCNA*) (**Fig 3d**) as well as genes involved in DNA damage response (e.g., *BARD1, RAD51C* and *H2AX*) (**Fig 3d**) and DNA repair (RECQL4, XPC and OGG1). Strong enrichment of genes for receptor tyrosine kinases (e.g., ERBB3, TRIB3 and GRB2) suggests a potential involvement SYCP1 in cancer signaling pathways. Additionally, we observed abundant SYCP1 binding at proximal promoters and/or gene bodies of approx. 30% of histone genes (45 genes out of total 118 in human geneome), what could suggest a possible role of SYCP1 in chromatin regulation.

**Figure 3.**
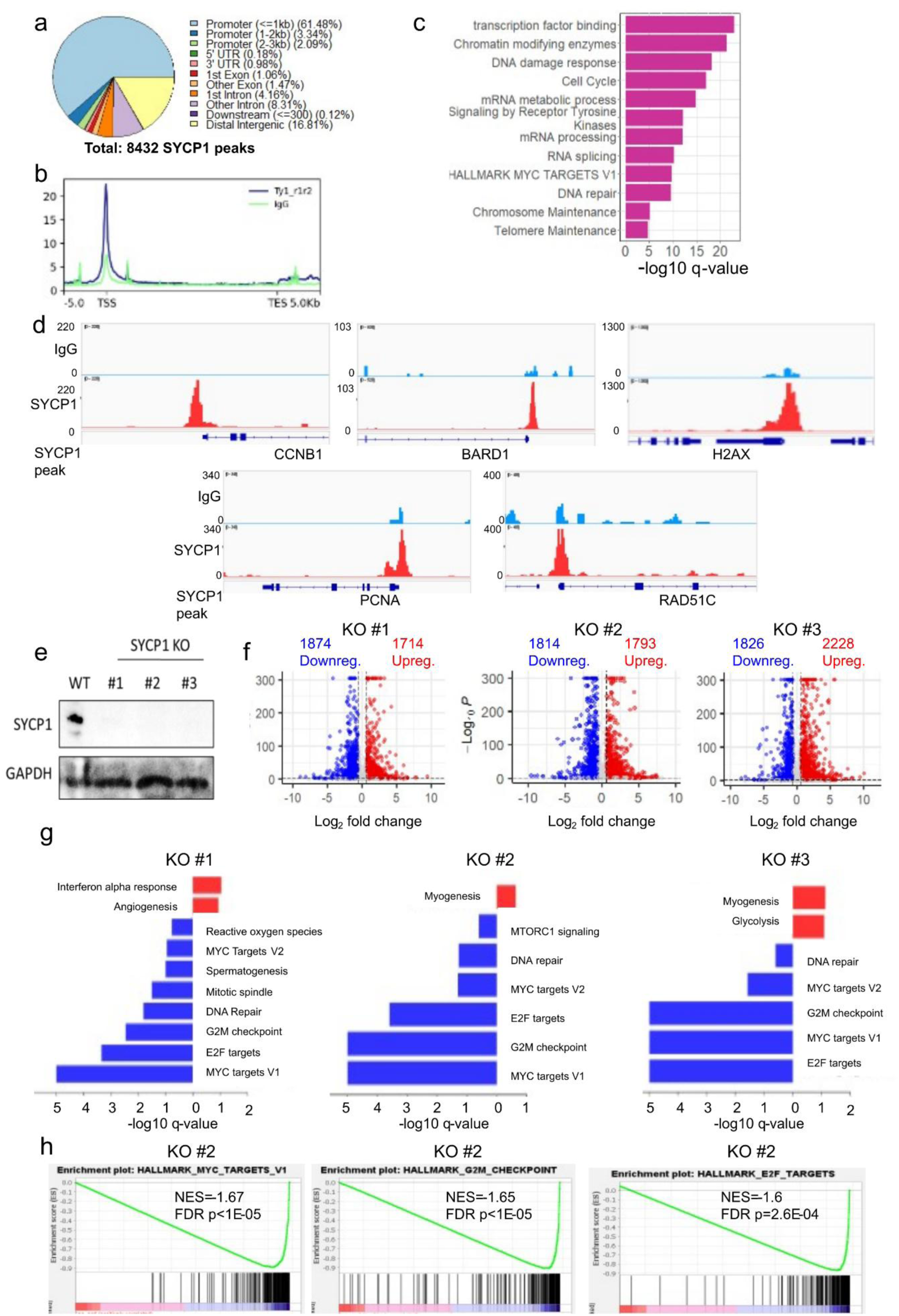
SYCP1 binds to the promoters of genes responsible for DNA damage response and cell cycle. (a) The total number and distribution of SYCP1 CUT&Tag DNA binding regions across genomic elements. (b) Metagene plot of SYCP1 binding levels within SYCP1-bound genes. The y axis represents SYCP1 CUT&Tag read density, compared to IgG negative control. (c) Pathway enrichment analysis of genes associated with SYCP1 bound regions (bar plot). (d) IGV browser view of selected CUT&Tag SYCP1 binding regions at the promoters of CCNB1, BARD1, H2AX, PCNA and RAD51C genes. (e) SYCP1 protein levels in WT and CRISPR KO MCF7 lines. (f) Volcano plots of differential gene expression analysis (RNA-seq data) across SYCP1 CRISPR KO cell lines, pathways enrichment analyses using Metascape software represented as two-directional bar plot (g) as well as using GSEA software (HALLMARK database) shown as GSEA enrichment score plot (h).

### SYCP1 regulates transcription of genes involved in cell cycle control and DNA damage response

To determine whether SYCP1 binding to gene promoters modulates transcriptional activity, we generated *SYCP1* knock-out (KO) MCF7 cell lines using CRISPR/Cas9-mediated genome editing. Three independent KO clones were selected based on confirmed absence of SYCP1 protein expression (**Fig. 3e**) and subjected to transcriptomic profiling by RNA-seq.

Differential gene expression (DGE) analysis revealed a highly reproducible transcriptomic signature across all three *SYCP1* KO clones relative to wild-type MCF7 cells (**Fig. 3f**). Notably, the most significantly downregulated pathways included cell cycle progression and DNA damage response (DDR) and repair mechanisms (**Fig. 3g–h**). Key downregulated genes included *CCNB1*, *CDC20*, *CDKN2C*, *CDKN2D*, *CENPA*, *H2AX*, *MCM4*, *MCM7*, *MYC*, *PCNA*, *POLD1*, *RAD51C*, *PLK4*, and *PRC1*—all of which are critical regulators of genomic maintenance and mitotic fidelity. In contrast, *SYCP1* loss led to transcriptional upregulation of genes associated with angiogenesis, glycolytic metabolism, and myogenic differentiation, such as *NFATC4*, *UVSSA*, *BMF*, *ATP6V1G1*, *TOM1*, and *ELAPOR1*.

To identify direct transcriptional targets of SYCP1, we integrated the transcriptomic data from the *SYCP1* KO clones (828 common downregulated and 781 common upregulated genes) with SYCP1 chromatin occupancy profiles generated by CUT&Tag. This integrative analysis identified 290 genes that were both significantly downregulated and had SYCP1 occupancy at their promoter regions (**Fig. 4a–b**). In contrast, only 170 promoter-bound genes were significantly upregulated upon *SYCP1* deletion. Hypergeometric testing revealed a significant overrepresentation of SYCP1-bound genes among the downregulated targets, whereas the overlap among upregulated genes was weakly under-enriched. These findings suggest that SYCP1 primarily acts as a transcriptional activator at the promoters it binds, directly regulating a gene network enriched in cell cycle, DDR, and RNA metabolic processes (**Fig. 4c**). Of particular interest, histone gene clusters were prominently enriched among the direct SYCP1 targets (**Suppl. Fig. 5**), implying a potential role for SYCP1 in chromatin assembly or replication-associated transcription.

**Figure 4.**
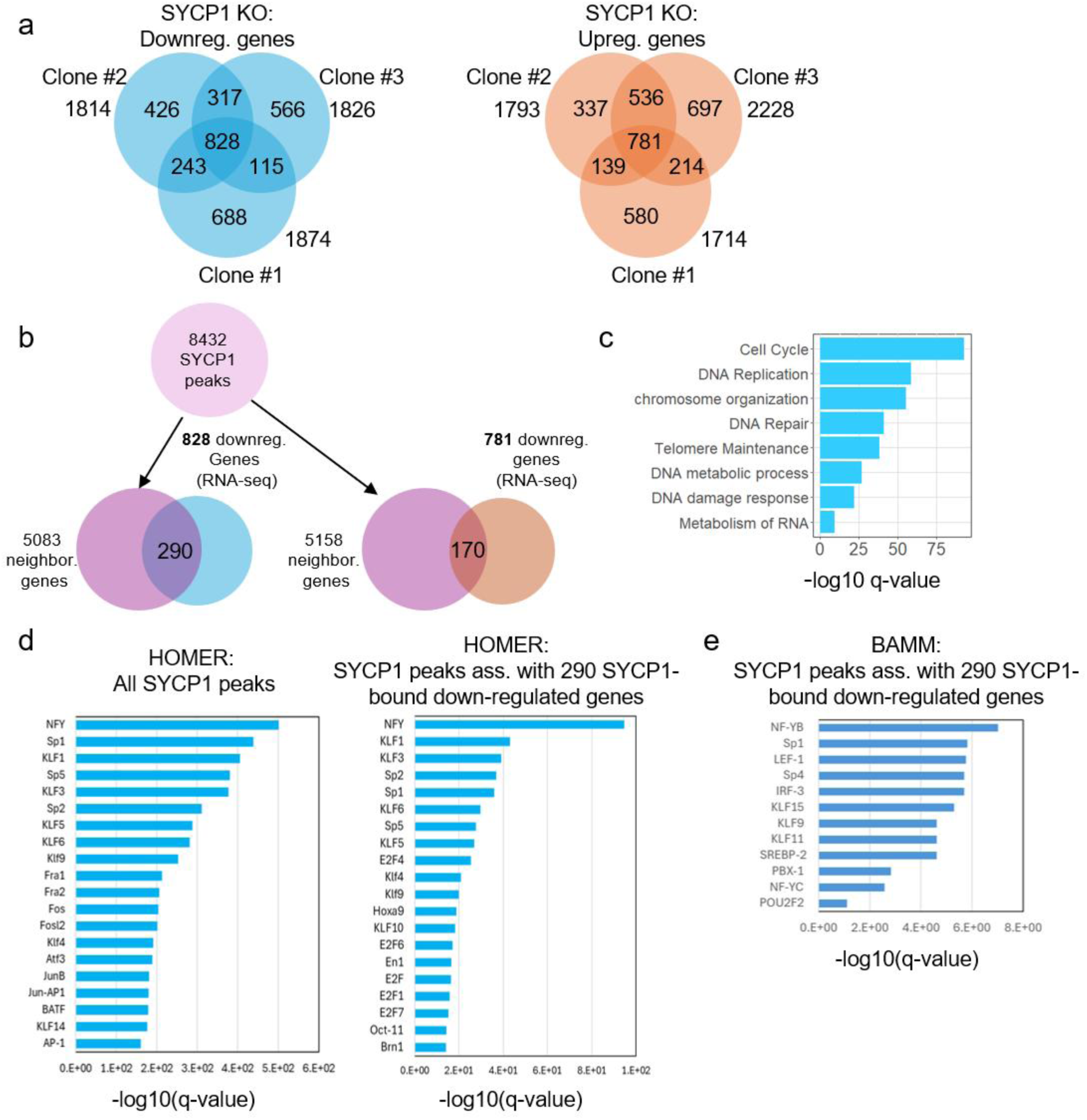
SYCP1 regulates expression of genes involved in DNA damage response and cell cycle. (**a-b**) Venn diagrams summarizing the results of RNA-seq analyses in three MCF7 SYCP1 KO cell lines; (**c**) Comparison of RNA-seq and CUT&Taq data analyses for SYCP1: the down-regulated overlapping gene list (290 genes) is strongly over-enriched (1.37-fold, p=2.2e-10) and the upregulated overlapping gene list (170 genes) is weakly under-enriched (1.18-fold, p=0.004), hypergeometric test; q-val=0.01 (**d**) Bar plot showing pathways enrichment analysis for all SYCP1 promoter bound genes. Transcription factor motif enrichment analysis using HOMER (http://homer.ucsd.edu/homer/), qval_= 1E-05 (**e**) Bar plot showing pathways enrichment analysis for 290 SYCP1 promoter bound and SYCP1 KO down-regulated genes. Transcription factor motif enrichment analysis using HOMER (http://homer.ucsd.edu/homer/), (**e**) and (**f**), BAMM software (https://bammmotif.mpibpc.mpg.de/) (**g**). Enriched motifs with correspondingly assigned TFs are shown at the bottom.

To further characterize the transcriptional regulatory landscape at SYCP1-bound loci, we performed transcription factor (TF) motif enrichment analysis within ±150 bp of SYCP1 peak summits. Using two distinct computational pipelines—HOMER and BAMM—we interrogated both the global SYCP1 binding landscape (**Fig. 4d**) and the subset of 290 direct target genes (**Fig. 4e–f**). Motifs enriched in these regions were annotated to known TFs using curated TF-binding databases. Both tools converged on a core cohort of transcription factors implicated in cell cycle regulation, chromatin remodeling, and DNA repair (**Fig. 4g**), indicating that SYCP1 may function cooperatively with established oncogenic TFs to control proliferation and genome maintenance programs.

Together, these results establish SYCP1 as a chromatin-associated regulator that directly activates genes essential for cancer cell proliferation and survival by modulating transcriptional programs governing the cell cycle and genome stability.

### SYCP1 promotes cancer cell proliferation and migration

Our transcriptomic and chromatin profiling analyses identified SYCP1 as a regulator of gene networks governing cell cycle progression and cell motility (**Figs. 3–4**). To directly assess the functional impact of SYCP1 depletion on cancer cell behavior, we first performed acute SYCP1 knockdown in MCF7 breast cancer cells. Loss of SYCP1 resulted in a rapid and profound proliferative arrest, with near-complete cessation of cell growth observed within 72 hours post-silencing (**Fig. 5a**). Similar growth inhibition phenotypes were observed in additional cancer models, underscoring a broader functional requirement for SYCP1 in tumor cells (**Suppl. Fig. 6**).

**Figure 5.**
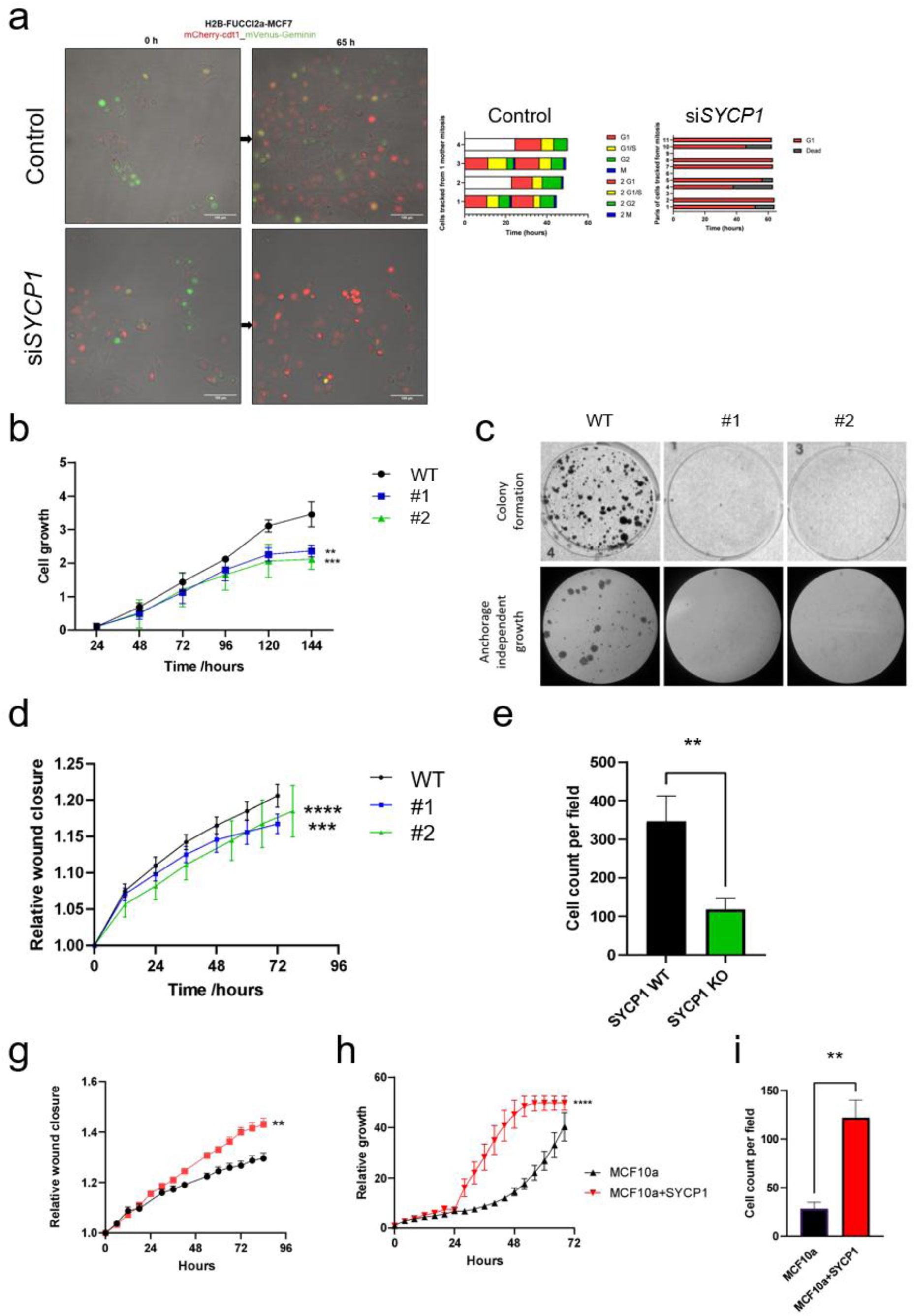
SYCP1 controls proliferation and migration of breast cancer cells. (**a**) MCF7 cells stably transfected with H2B-Fucci2a were transfected with control (SCR), SYCP1 targeting siRNAs or a mock transfection. 24 hours post transfection, live cell imaging was initiated, with images acquired every 15 minutes for 65 hours. Images show fields of view at 0 and 65 hours, for Mock and siSYCP1_B transfections. Stacked bar graphs show cell cycles from pairs of individual cells that were analysed by tracking daughters of a mitosis, either through two complete cell cycles or for the remaining duration of the timelapse. (**b**) WT and SYCP1 CRISPR KO lines of MCF7 cells were seeded at equal density and cell number were tracked with Incucyte live cell imaging for 90hrs. (**c**) WT and SYCP1 CRISPR KO lines of MCF7 cells were seeded at equal density on standard plates (colony formation) or in soft agar (anchorage independent growth) and number of colonies after a week was compared. (**d**) WT and SYCP1 CRISPR KO lines of MCF7 cells were seeded at equal density and scratched to create a wound with closure monitored by Incucyte life cell images for 90hrs. (**e**) WT and SYCP1 CRISPR KO lines of MCF7 cells were seeded at equal density into transwells in serum starved media with serum rich media at the other side of the chamber. Following 12 hours incubation cells that migrated through the transwell were counted. (**f**)WT and SYCP1 transfected MCF10A cells were seeded at equal density and cell number were tracked with Incucyte live cell imaging for 90hrs. (**g**)WT and SYCP1 transfected MCF10A cells were seeded at equal density and scratched to create a wound with closure monitored by Incucyte live cell images for 90hrs. (**i**) WT and SYCP1 transfected MCF10A cells were seeded at equal density into transwells in serum starved media with serum rich media at the other side of the chamber. Following 24 hours incubation cells that migrated through the transwell were counted.

We next examined the long-term effects of complete SYCP1 ablation by analyzing stable *SYCP1* CRISPR knock-out (KO) clones. The markedly reduced efficiency in isolating viable KO clones highlighted a dependency of MCF7 cells on SYCP1 for sustained proliferation. Once established, *SYCP1*-null clones exhibited significantly impaired proliferative capacity (**Fig. 5b**), diminished clonogenic potential (**Fig. 5c**), and reduced migratory behavior, as assessed by both transwell and scratch-wound assays (**Fig. 5d–e**).

To determine whether SYCP1 expression alone is sufficient to induce a transformed phenotype, we ectopically expressed SYCP1 in non-tumorigenic MCF10A breast epithelial cells. SYCP1 expression drove a dramatic increase in cell proliferation, with a >20-fold enhancement in viable cell number relative to vector control (**Fig. 5f**). Similarly, SYCP1-expressing MCF10A cells displayed enhanced migratory potential in both directed and collective migration assays (**Fig. 5g–h**). Together, these findings reveal that SYCP1 is both necessary and sufficient to drive key tumorigenic properties, including uncontrolled proliferation and enhanced motility.

### DNA-binding of SYCP1 is critical for its function

SYCP1 is a 976 amino-acid coiled-coil protein that mediates synapsis between homologous chromosomes during meiosis. This is achieved through the C-termini binding to chromosomal DNA, with the N-terminal ends of juxtaposed SYCP1 molecules interacting head-to-head in a zipper-like manner midway between synapsing chromosomes [11, 22]. Hence, we wondered whether the pro-proliferative effect of SYCP1 is due to its DNA-binding activity and/or its head-to-head self-assembly (**Fig 6a**). To test this, we analysed a C-terminal construct (amino-acids 640-976) that is dimeric as it contains the end of the coiled-coil, includes the full DNA-binding region, and was previous shown to bind to DNA *in vitro* [22]. Upon over-expression, SYCP1 640-976 had localised to chromatin (**Fig 6b,c**) and stimulated growth in knockout MCF7 cells at a level comparable to the full-length molecule (**Fig 6d**). Thus, we conclude that the pro-proliferative effect of SYCP1 is mediated solely by its DNA-binding region and does not depend on self-assembly of its N-termini.

**Figure 6.**
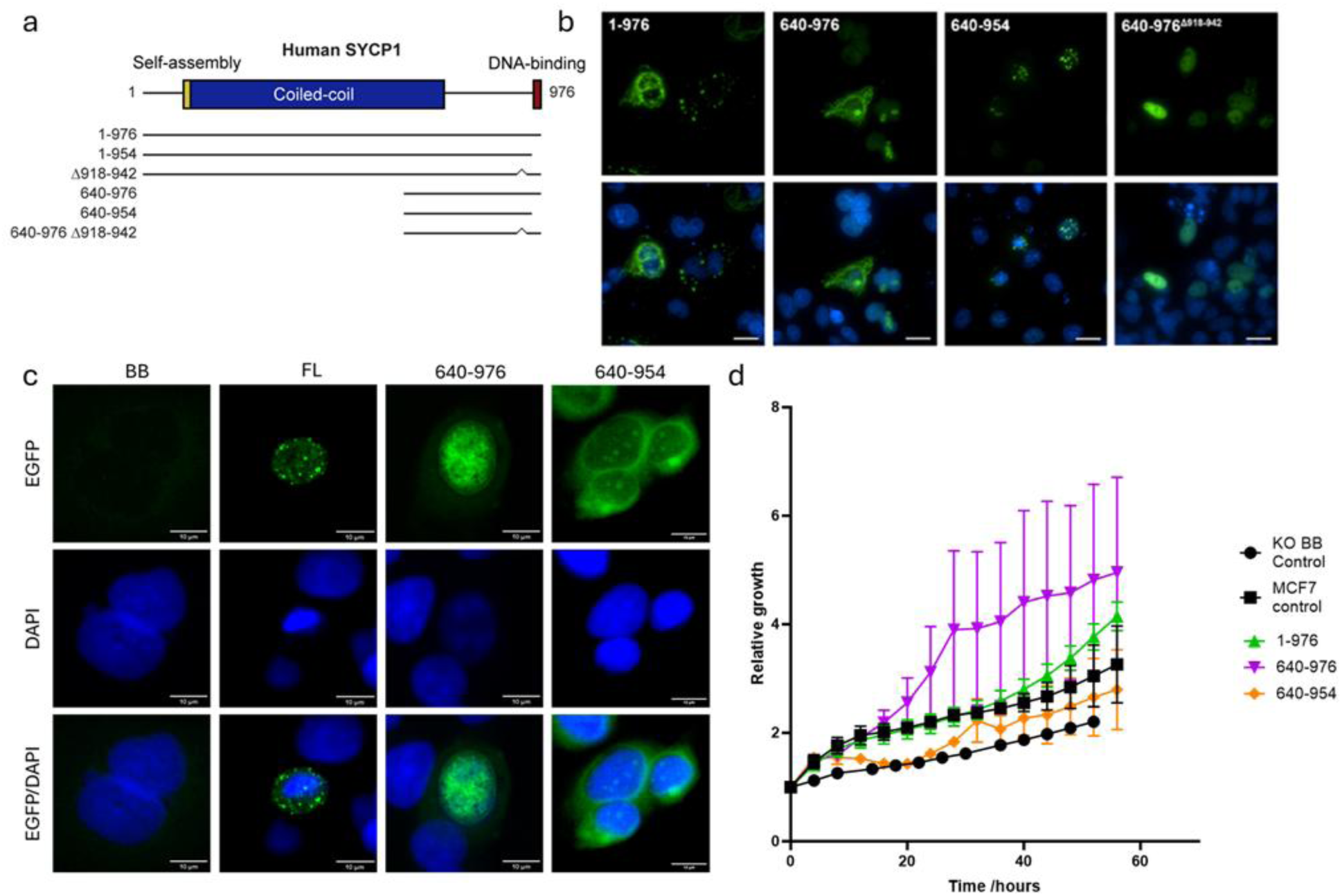
DNA-binding of SYCP1 is critical for its function. **(a)** Structure of human SYCP1 and mutants used in this study. (b) Transient overexpression of EGFP-SYCP1 constructs in COS7 cells. The numbers correspond with the construct boundaries of the amino acid sequence. 20 µm scalebar. (c) SYCP1 truncations were investigated in MCF7 SYCP1 KO cells. (d) MCF7 SYCP1 KO cells were transfected with full length, and truncated SYCP1 and tracked with incucyte live imaging for 90 hours.

The SYCP1 640-976 construct contains a dimeric coiled-coil (amino-acids 640-783), followed by predicted unstructured sequence, which includes substantial basic charge at its C-terminal end that likely mediates, or contributes to, its DNA-binding (**Fig 6a**). Accordingly, we found that deletion of the last 22 amino-acids of the protein (in construct 640-954) abrogated the stimulation of growth (**Fig 6d**), whilst retaining nuclear localisation (**Fig 6b, c**). Hence, DNA-binding mediated by basic amino-acids at the C-terminal tip of the protein is required for SYCP1’s pro-proliferative effect. We also identified an internal deletion with SYCP1’s unstructured C-terminus (Δ918-942) that had a similar effect in abrogating growth stimulation, whilst retaining nuclear localisation (**Fig 6b**). Importantly, this deletion does not include overt DNA-binding basic patches, indicating that the wider structure of the SYCP1 C-terminus is likely important for its pro-proliferative effect.

Having demonstrated these effects using the C-terminal 640-976 construct, we next assessed the same deletions within full-length SYCP1. We found that SYCP1 1-954 truncation had decreased nuclear localisation, while del918-942 variant was largely unable to localise to the nucleus (**Fig 7a,b**). Hence, the two regions that are necessary for promoting growth also contribute to nuclear localisation, and loss of either sequence mislocalises SYCP1. We performed rescue experiments in MCF7 *SYCP1* KO cells and found that while full length SYCP1 was able to rescue all cellular function to WT levels variants with nuclear localisation defects were not able to restore the cell cycle profile (**Fig 7c**), proliferation (**Fig 7d**) or migration (**Fig 7e,f**). The same relationship was observed when SYCP1 variants were introduced to MCF10a cells (**Fig 7g-j**).

**Figure 7.**
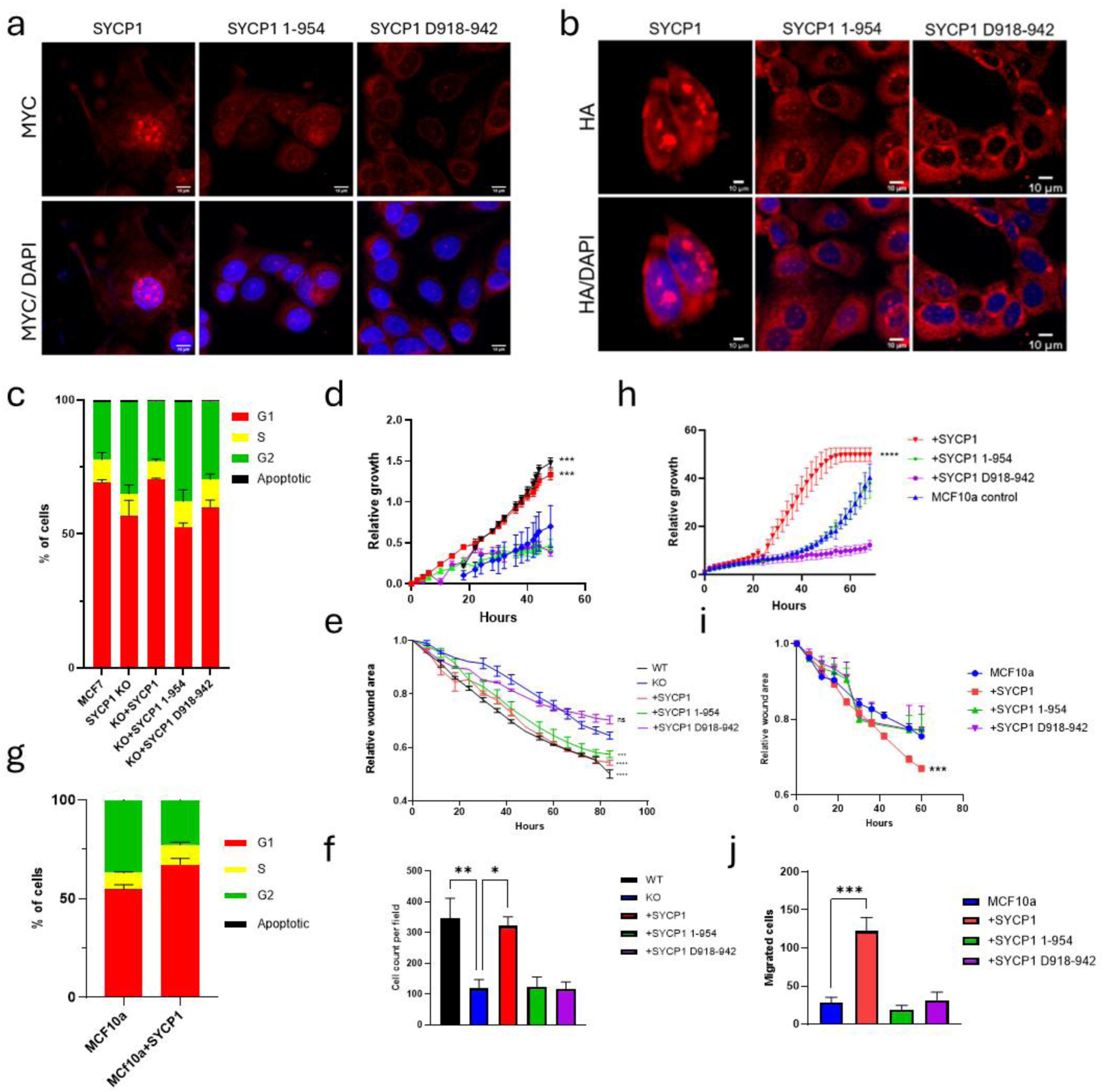
Nuclear localisation of SYCP1 is critical for pro-proliferative and migratory function. (**a-b**) SYCP1 truncation 1-954 and deletion D918-942 were investigated in MCF7 cells after identification as possible NLS’s. Nuclear localisation was completely lost in D918-942 and a reduction was observed in SYCP1 1-954, for both the N-term MYC tagged SYCP1 (**a**) and the C-term HA tagged (**b**). (**c**) Cell cycle function was investigated through the use of guava flow, the shorter G1 observed in KO cells was rescued only by the nuclear localised full length SYCP1. (**d-e**) MCF7 SYCP1 KO cells were transfected with full length, 1-954 and D918-942 SYCP1 and tracked with incucyte live imaging for 90 hours, only full length rescued proliferation (**d**) and wound closure rate (**e**). (**f**) Cells were seeded at equal density into transwells in serum-starved media with complete media at the other side of the chamber. After 24 hours of incubation, migrated cells were counted, only full length SYCP1 rescued KO cells to WT levels of migration. MCF10a WT and SYCP1 transfected cells were subjected to cell cycle analysis (**g**). WT MCF10a cells and cells transfected with SYCP1 variants were seeded at equal density for 90 hours and both proliferation (**h**) and wound closure (**i**) rate were determined. (**j**) WT MCF10a cells and cells transfected with SYCP1 variants were seeded at equal density into transwells in serum-starved media with complete media at the other side of the chamber. After 24 hours, cells that migrated through the transwell were counted, only full length SYCP1 increased MCF10a cells migration rate.

### SYCP1 is a component of promoter-specific DNA-binding complexes that modulate transcription factor activity

To elucidate the molecular mechanisms by which SYCP1 regulates gene expression in cancer cells, we employed proximity-dependent biotin labeling (BioID2) to define the SYCP1 interactome. Mass spectrometry analysis revealed that SYCP1 associates with chromatin-bound proteins, including core histones and chromatin-modifying enzymes. Notably, SYCP1 was enriched in complexes involved in promoter-specific DNA binding, a pattern that was corroborated by SYCP1 CUT&Tag profiling, which revealed co-localization with gene promoters and enriched signal over single-stranded and oxidized DNA regions (**Fig. 8a**, i–ii). SYCP1 was also found in proximity to histone methyltransferases and DNA endonucleases, suggesting a broader role in chromatin state modulation.

**Figure 8.**
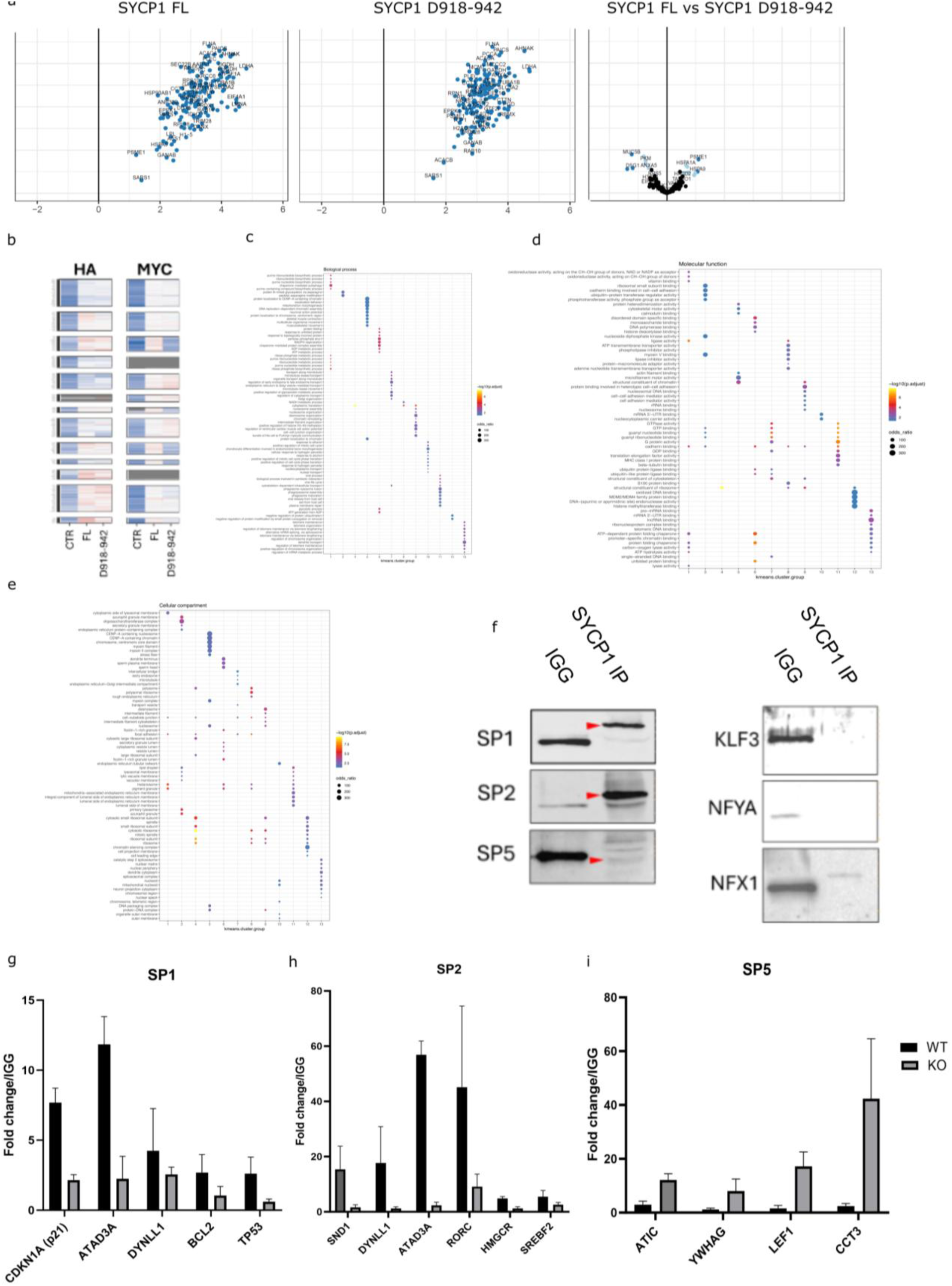
BIOID2 interactome of WT SYCP1 vs D918-942. MCF7 cells with stable integration of BIOID2-SYCP1 or BIOID2-SYCP1 D918-942 were biotinylated for 24 hours. Pulldowns were sent for mass spectrometry TMT analysis. (**a**) Detected proteins were compared to control in SYCP1 (**i**) and SYCP1 D918-942 (**ii**) as well as to each other (**iii**). (**b**) 13 clusters were detected based on expression patterns between the control, full length and truncated version tagged at both the N and C terminus. Enrichment of pathways was detected in each cluster showing differences in cellular component analysis (**c**), biological processes (**d**) and molecular function (**e**). (**f**) Western blot of SP1, SP2, SP5, KLF3, NFYA and NFX1 in an immunoprecipitation of SYCP1 in MCF7 cells. (**g**) SP1 ChIP with qPCR of SYCP1 target genes including shared SYCP1 and SP1 targets ATAD3A, p21 and DYNLL1. Data was normalised to IgG control and represents 3 biologically independent repeats, error bars show SEM. (**h**) SP2 ChIP with qPCR of SYCP1 target genes including shared SYCP1 and SP2 targets SIND1, DYNLL1 and ATAD3A. Data was normalised to IgG control and represents 3 biologically independent repeats, error bars show SEM. (**i**) SP5 ChIP with qPCR of SYCP1 target genes including shared SYCP1 and SP5 targets ATIC and YWHAG. Data was normalised to IgG control and represents 3 biologically independent repeats, error bars show SEM.

To probe the functional relevance of SYCP1’s C-terminal region, we compared the interactomes of wild-type SYCP1 and a truncated mutant lacking amino acids 918–942 (Δ918–942), which fails to rescue SYCP1-dependent phenotypes in cancer cells. BioID2-tagged SYCP1 constructs (WT and Δ918–942) and empty vector controls were stably expressed in MCF7 cells, biotinylated for 24 hours, followed by streptavidin enrichment and tandem mass tag (TMT)-based quantitative proteomics (**Fig. 8a**, iii). Comparative analysis revealed distinct interaction profiles between WT and mutant SYCP1, with 13 discrete protein clusters showing differential proximity (**Fig. 8b**). The Δ918–942 variant lost binding to nuclear proteins and, critically, failed to interact with mitotic cell cycle regulators and components of adhesion complexes including cadherin-associated proteins, focal adhesion machinery, and cell–cell junctional proteins (**Fig. 8c–e**).

Integrating SYCP1’s BioID2 interactome with CUT&Tag co-occupancy data, we prioritized a set of transcription factors (TFs) that are likely to co-function with SYCP1 at chromatin. These included SP1, SP2, SP5, KLF3, NFYA, and NFX1. Co-immunoprecipitation experiments in MCF7 cells demonstrated that SYCP1 selectively binds to SP1, SP2, and SP5, but not to KLF3, NFYA, or NFX1 (**Fig. 8f**).

To test whether SYCP1 regulates the chromatin association of SP-family TFs, we examined SP1, SP2, and SP5 occupancy at shared target genes upon SYCP1 loss (**Fig. 8g–i**). ChIP-qPCR revealed that SYCP1 depletion impaired SP1 and SP2 binding at co-targeted promoters (**Fig. 8g–h**), while SP5 occupancy was conversely increased (**Fig. 8i**), suggesting a competitive or compensatory regulatory relationship. These findings indicate that SYCP1 modulates promoter accessibility and transcriptional output through specific interaction with and regulation of SP-family transcription factors, positioning SYCP1 as an integral component of oncogenic gene regulatory circuits in cancer cells.

### SYCP1 modulates chemotherapy response

In its canonical meiotic role, SYCP1 is a core component of the synaptonemal complex (SC), which bridges homologous chromosomes and facilitates the repair of programmed double-strand breaks (DSBs) during meiosis [11]. Our findings in cancer models revealed that SYCP1 is aberrantly expressed in the nucleus, binds chromatin, and regulates transcription of DNA damage response and repair genes (**Figs. 2–4**). Given this context, we hypothesized that SYCP1 might be functionally responsive to genotoxic stress and directly localize to DNA damage sites.

Given that patients with high SYCP1 expression exhibit significantly shorter overall and relapse-free survival across multiple cancer types (**Sup Fig. 1**), we investigated whether SYCP1 contributes to chemoresistance by facilitating DNA repair. We tested this hypothesis using the chemotherapeutics gemcitabine and cisplatin, a combination frequently used in neoadjuvant therapy for non-operable breast cancer [23]. SYCP1 knockout sensitized MCF7 breast cancer cells to both drugs (**Fig. 9a**), while having no impact on the response to rucaparib (a PARP1 inhibitor) or MYCi361 (a MYC pathway inhibitor). A similar drug response profile was observed in non-transformed MCF10A cells ectopically expressing SYCP1, compared to untransfected controls (**Fig. 9b**), suggesting that SYCP1 enhances resistance to genotoxic stress by modulating early stages of the DNA repair cascade.

**Figure 9.**
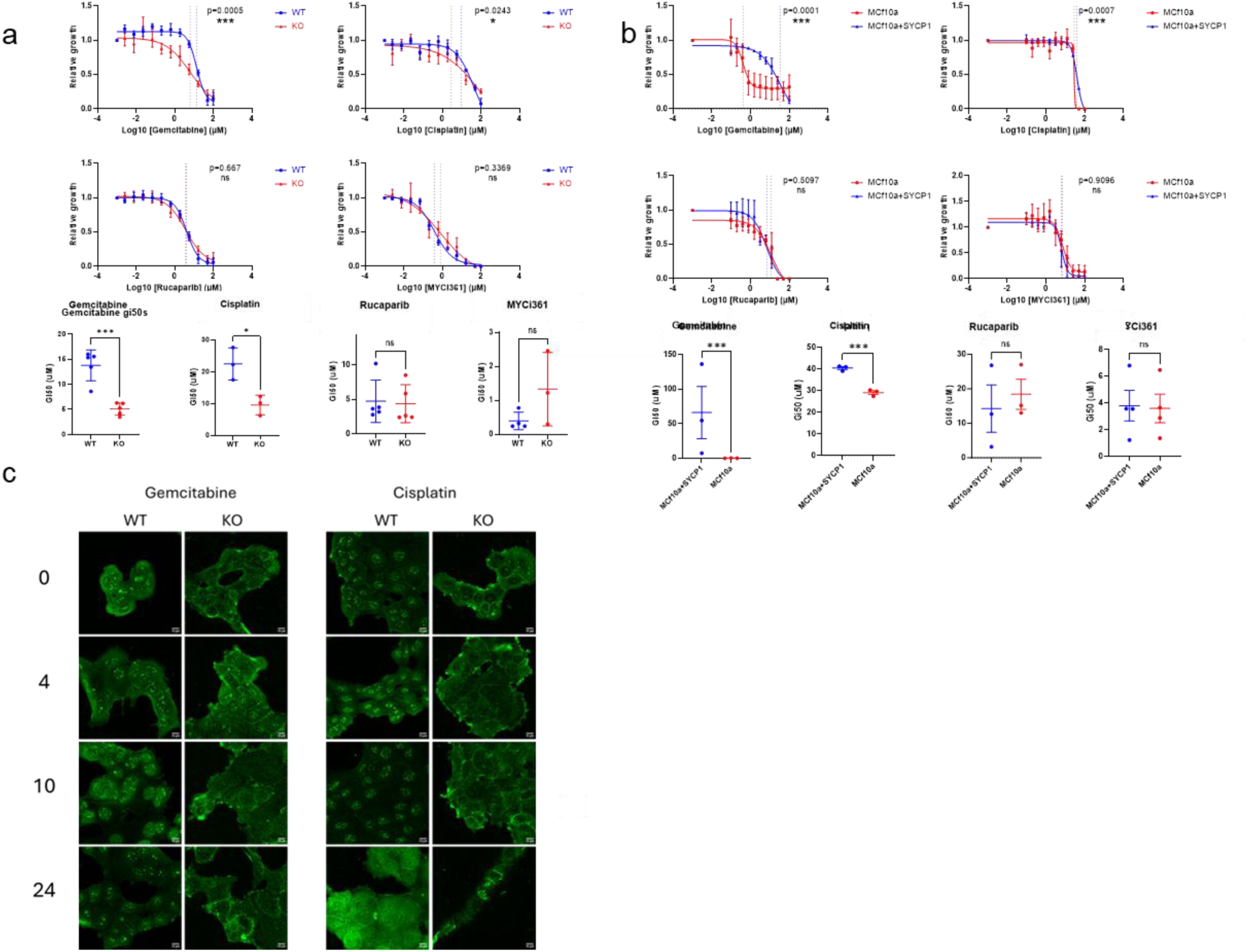
SYCP1 promotes resistance to chemotherapy in breast cancer cells. (**a**) Gi50s of WT and SYCP1 KO MCF7 cells in response to gemcitabine, cisplatin, rucaparib and MYCi361. (**b**) Gi50s of MCF10a cells and MCF10a transfected with SYCP1 in response to gemcitabine, cisplatin, rucaparib and MYCi361. (**d**) DNA damage marker gH2ax in MCF7 WT or KO cells, at 0 hours (damage induction) and after 4, 10 or 24 hours recovery, in response to gemcitabine or cisplatin.

To further explore SYCP1’s function in DNA repair kinetics, we examined the formation and resolution of γH2AX foci following chemotherapeutic insult (**Fig. 9c**). In wild-type cells, gemcitabine and cisplatin treatment induced pronounced γH2AX foci that persisted for up to 24 hours. In contrast, SYCP1-deficient cells formed fewer foci with more rapid resolution, suggesting that SYCP1 may contribute to foci formation and stabilization of the damage response.

Together, these data position SYCP1 as a regulator of the DNA damage response in cancer cells, facilitating both recruitment to DNA lesions and influencing repair kinetics—mechanistic features that may underlie its association with poor clinical outcomes and chemoresistance.

### SYCP1 hinders DNA damage repair in breast cancer patients and correlates with poor prognosis

To investigate the clinical relevance of SYCP1 re-expression, we analyzed a large cohort of breast cancer patients, focusing on protein-level expression due to the previously noted discordance between SYCP1 mRNA and protein abundance (**Fig. 1**). Immunohistochemical analysis revealed that SYCP1 protein was aberrantly expressed in the vast majority of patient samples (**Fig. 10a**). SYCP1 levels positively correlated with RAD51 expression (**Fig. 10b**). These observations support a role for SYCP1 at sites of DNA damage in vivo, consistent with our *in vitro* findings.

**Figure 10.**
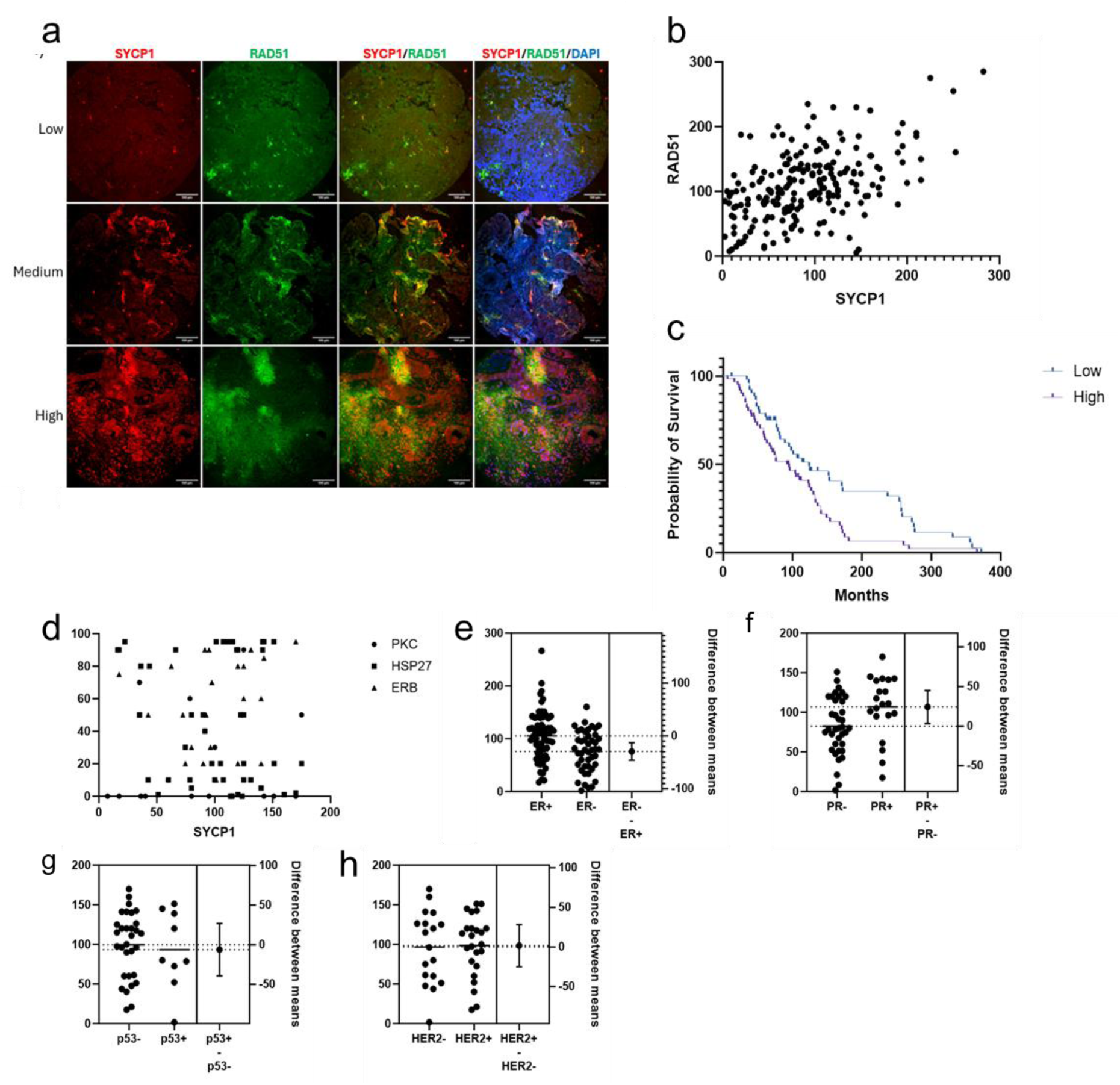
SYCP1 hinders DNA damage repair in breast cancer patients. (**a**) TMAs from breast cancer patients were stained for both RAD51 and SYCP1, staining levels vary between patients. (**b**) RAD51 and SYCP1 have a positive correlation in the cohort. p<0.0001, r^2^=0.2438. (**c**) Kaplan meier of the impact of SYCP1 on cohorts survival. Gehan-Breslow-Wilcoxon test p=0.0191 Median survival for SYCP1 Low 124.0, SYCP1 High 95.00, Ratio (and its reciprocal) 1.305, 0.7661, 95% CI of ratio 0.8721 to 1.954; 0.5119 to 1.147, n=114. (**d**) No correlation between SYCP1 and PKC, HSP27 or ERB. Both ER+ (**e**) and PR+ (**f**) patients had higher histoscores. Histoscores remained the same despite P53 (**g**) or HER2 (**h**) patient status.

Similar corelation patterns of SYCP1 and RAD51 were observed in additional cancer types, including sarcoma, colon, kidney, and liver cancers (**Sup Fig. 7**). Notably, SYCP1 expression did not correlate with the expression of unrelated stress markers such as HSP27 or PKC (**Fig. 10d**), indicating specificity in its correlation with DNA repair machinery.

We next examined whether SYCP1 expression was associated with clinical features in breast cancer. Estrogen receptor–positive (ER+) patients exhibited significantly higher SYCP1 protein levels compared to ER– patients (**Fig. 10e**), a trend also observed for progesterone receptor (PR) status (**Fig. 10f**). However, SYCP1 levels were not significantly associated with TP53 mutational status or homologous recombination (HR) proficiency (**Fig. 10g,h**), suggesting that SYCP1 expression is not merely a surrogate marker of global genomic instability.

Critically, high SYCP1 expression was significantly associated with reduced overall survival in breast cancer patients (**Fig. 10c**), reinforcing its potential as a clinically relevant marker of poor prognosis. These findings suggest that aberrant expression of SYCP1 in cancer may impair DNA repair fidelity, promoting genome instability and therapeutic resistance.

## Discussion

Our study uncovers an unexpected oncogenic role for the meiotic protein SYCP1 in somatic cancer cells, where it regulates transcriptional programs, influences DNA damage repair, and promotes tumor progression (**Fig. 11**). Classically defined as a structural component of the synaptonemal complex essential for meiotic homologous recombination [10], SYCP1 is aberrantly re-expressed in cancer and repurposed for non-canonical functions in genome maintenance. These findings underscore a broader paradigm in tumor biology, cancer cells exploit developmental pathways, co-opting germline proteins to overcome genomic instability and therapeutic pressure.

**Figure 11.**
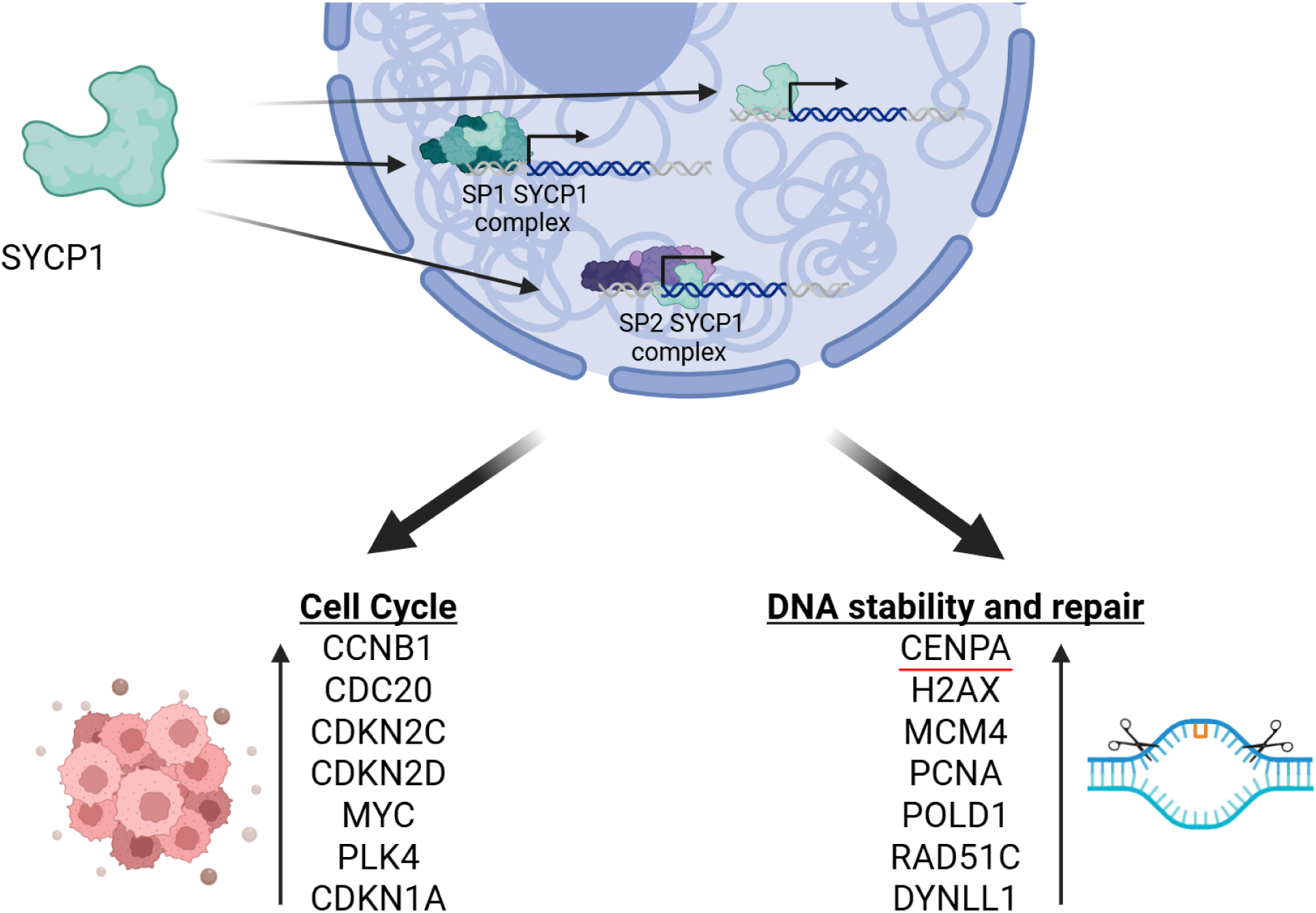
Proposed model of SYCP1 function in cancer cells. In cancer cells, the normally meiotic protein SYCP1 is aberrantly expressed and translocates to the nucleus, where it adopts a non-canonical regulatory role. Once nuclear, SYCP1 associates with the promoters of a distinct subset of genes, including cell cycle and proliferation-related targets, via interactions with the transcription factors SP1 and SP2. This complex promotes transcriptional activation, leading to increased expression of oncogenic programs. SYCP1-driven gene expression changes are associated with enhanced proliferative capacity, resistance to DNA damage, and disruption of normal chromatin architecture. Our model suggests that SYCP1 acts as a chromatin-bound transcriptional co-regulator in cancer, repurposed from its canonical structural role in meiosis.

### Moonlighting Functions of SYCP1 Beyond Structural Support

In meiosis, SYCP1 facilitates chromosome synapsis by bridging homologous chromosomes [22]. In contrast, in cancer cells, we show that SYCP1 localizes to chromatin, where it regulates gene expression programs associated with cell cycle control and DNA repair. Mechanistically, SYCP1 modulates chromatin recruitment of SP-family transcription factors, particularly SP1 and SP2, which are key regulators of TATA-less housekeeping genes. SP1 is a well-established oncogenic TF that promotes proliferation in various malignancies, including breast cancer [24], and correlates with poor clinical outcomes [25]. SP2 similarly contributes to tumor formation [26]. Conversely, SP5 has more limited expression and has been linked to pro-apoptotic signaling, including as a WNT pathway target [27]. We propose that SYCP1 promotes tumorigenesis by enhancing SP1 and SP2 recruitment to chromatin while suppressing SP5 activity, collectively driving cell cycle progression and migration.

Our proximity labeling and CUT&Tag experiments reveal that SYCP1’s interactome is context-dependent, shifting from structural synapsis-associated partners in meiosis to chromatin regulators and transcription factors in cancer. This functional plasticity mirrors findings for other meiotic proteins aberrantly re-expressed in tumors. For instance, SPO11 and HORMAD1—essential for meiotic recombination—have been shown to disrupt DNA repair when expressed in somatic cells [4, 28, 29].

### SYCP1 and DNA Damage Repair: A Double-Edged Sword?

Our data show that SYCP1 regulates the transcription of key DNA repair genes, including BARD1, RAD51C, and H2AX. Loss of SYCP1 sensitizes cancer cells to DNA-damaging agents such as cisplatin and gemcitabine, indicating a role in promoting repair. However, rather than supporting high-fidelity repair, SYCP1 may facilitate a non-canonical, potentially error-prone pathway that enables cancer cell survival. This is consistent with the idea that tumors often rely on alternative repair strategies to maintain viability under genotoxic stress.

Given the association between defective repair, chromosomal aberrations, and aggressive tumor behavior, further mechanistic studies are warranted to elucidate whether SYCP1 contributes to mutagenic repair outcomes that drive clonal evolution and resistance.

## Materials and Methods

### Cell culture, transfections

MCF7 breast cancer cells were obtained from the American Type Culture Collection (Manassas, USA). Cell lines were maintained in DMEM media supplemented with 2 mM l-glutamine (Invitrogen) and 10% (v/v) fetal calf serum (FCS) at 37 °C in 5% CO_2_. Cell lines were never maintained for more than 30 passages or 2 months of continuous culturing.

Cell lines were tested for mycoplasma on a tri-monthly basis. Proliferation was measured by live cell imaging with the Incucyte system every 6 h for 156 h post-treatment. Transfections were performed using Lipofectamine2000 reagent (Invitrogen) following the manufacturer’s instructions.

### Generation of a doxycycline-inducible SYCP1-expressing MCF7 line

SYCP1 was overexpressed in MCF7 cells using the pCW-SYCP1_3xTy1 vector. The vector was transfected into MCF7 cells, and a stable clone established. SYCP1 expression was induced with 5 µg/mL doxycycline (dox).

### siRNA gene silencing and gene expression analysis

The *SYCP1* targeting siRNA sequences were (A) GACUCUGAUUUGGAGAAUUCA[dTdT], (B) GAGUUUCCAUUUGCAAAGACU[dTdT] and (C) GUCACUAUCAGGAAGGACU[dTdT] (Sigma). Cells were reverse transfected with siRNA using RNAiMax (Invitrogen) according to manufacturer’s instructions and incubated in culture media for 96 h prior to cell lysis and analysis. For real-time qPCR, extraction of RNA was carried out using Trizol (Invitrogen, cat no. 15596026) following the manufacturer’s instructions. One microgram of RNA was incubated at 65°C for 5 minutes, followed by incubation at 37°C for 2 minutes. cDNA was synthesised using M-MLV Reverse Transcriptase (Promega, cat no. M1701). For qPCR, we used SYBR Green PCR Master Mix (Invitrogen, cat no. 4309155), and StepOne systems (Applied Biosciences, cat no. 4376357), each sample was normalised to the *HPRT1* housekeeping gene. Three biological replicates were carried out for each reaction. Fold changes were calculated using the 2-ΔΔCT relative quantification method. (Livak and Schmittgen, 2001). Data was tested for parametric distribution. Parametric data was analysed using appropriate *t*-tests or ANOVA with Bonferroni’s comparison test for multiple group comparisons. Non-parametric data was analysed using Wilcoxon signed-rank test. By convention, \*\*\**p*-values < 0.001, \*\**p*-values < 0.01 and \**p*-values < 0.05.

### Genotyping CRISPR KO clones

SCYP1 knockout was generated through CRISPR/Cas9-mediated genome editing in MCF7 cells and was carried out by Synthego, using a guide RNA (gRNA) targeting exon 2 (5’ -AAGCAGCAGTCAGGTGTCTG-3’). Single clones were genotyped using Zero Blunt™ TOPO™ PCR Cloning Kit (Invitrogen, cat. no. 450245) according to the manufacturer’s instructions. Upon bacterial transformation, colonies were selected for plasmid isolation using the ZymoPURE Plasmid Miniprep Kit (Zymo Research, cat. no. D4214) and subsequently subjected to Sanger sequencing to identify editing events.

### Immunofluorescence

For Immunofluorescence, cells were fixed for 10 min in ice cold methanol followed by blocking in 1% BSA, staining overnight at 4C in primary antibodies then for 1 h at room temperature with goat anti-rabbit Alexa Flour594 (A32740) or Alexa Flour647 (A-21245). and rabbit anti-mouse Alexa Fluor488 (A-11059) Life Technologies secondary antibodies and DAPI. Mounting was performed in prolong antifade gold reagent (Life Technologies P36961) or Vectashield anti-fade medium (H-1000-10), imaging using LSM 880 confocal microscope and processed with Zen 2.6 blue edition software.

### Immunohistofluorescence

Tissue microarrays (TMA) containing 0.6 mm cores of breast cancer and control tissues including sarcoma, liver, colon and kidney cancer from retrospective cohorts were used. Our cohort included patients treated within the Merseyside Hospital Trust in the 90s and 2000s. Antigens were retrieved by microwaving the slides in 10 mM citrate pH 6.0 for 15 min followed by immunofluorescence. Samples were scored blind using the histoscore methodology ^30^. Briefly, percentage and intensity of staining for positive cells was estimated (0, 1, 2, 3) using the following equation: *H*-score = (% of cells with low level positivity) + 2 × (% of cells with medium level positivity) + 3 × (% of cells with high level positivity). For survival analysis, high marker levels were defined as a value in the third and fourth upper quarter of the population. Tissues were imaged at Liverpool University Centre for Cell Imaging (CCI).

### Chromatin fractionation

Cells were washed with PBS and trypsinised and pelleted. The cell pellets were washed with ice-cold PBS and lysed with CSK buffer (10 mM PIPES pH 7.0, 200 mM NaCl, 300 mM sucrose, 3 mM MgCl2, 1 mM DTT, 1 mM aprotinin, 1 mM pepstatin, 1 mM leupeptin, 2 mM PMSF) containing 0.5% (v/v) Triton X-100, and incubated on ice for 10 minutes. The lysate was then spun down at 1500 × g for 5 minutes at 4°C. The supernatant, was spun down at maximum speed for 30 minutes at 4°C. The pellet (chromatin fraction) was washed twice with CSK buffer containing 0.1% (v/v) Triton X-100. The pellet was resuspended in buffer C (20 mM Tris pH 7.9, 1.5 mM MgCl2, 420 mM KCl, 1 mM DTT, 1 mM aprotinin, 1 mM pepstatin, 1 mM leupeptin, 2 mM PMSF), and incubated in the cold room while rotating for 90 minutes, followed by five cycles of sonication on high setting. The chromatin fraction was then spun down at maximum speed for 30 minutes at 4°C. The supernatant was collected for downstream Western blotting.

### Colony formation assay

Cells were seeded in a 6-well plate at a density of 1,000 cells per well, left for 21 days then cells were washed with PBS and stained with crystal violet overnight.

### Transwell migration assay

Cells were seeded in the Transwell inserts in serum-free media and incubated for 24 hours. After incubation, the cells were washed once with PBS and fixed with ice-cold 75% (v/v) ethanol for 10 minutes, followed by additional washing with PBS. Crystal violet was added for overnight staining. Transwell inserts were then imaged using a microscope, and representative images were selected and processed using ImageJ.

### Anchorage-independent growth

Cells were plated into media containing 0.56% methylcellulose in a 24-well plate and incubated at 37 °C in 5% CO_2_ for 14 days followed by imaging with IncuCyte and automated colony counting.

### MCF7-H2B-Fucci2a live cell cycle analysis

The PiggyBac compatible pB-H2B-Cerulean-Fucci2a plasmid ^31^ was a kind gift from Richard Mort (Lancaster University). pB-H2B-Cerulean-Fucci2a and pbHybase, were transfected into MCF7 cells using Genejuice (70967-3, Merck). Stable transfectants were selected with puromycin and individual clones grown.

For live cell imaging, MCF7-H2B-Fucci2a cells were seeded onto glass-bottomed 24-well plates (P24-1.5P, Cellvis) and reverse transfected (Lipofectamine RNAiMax) with 10nM Control or siSYCP1_B siRNAs. 24 hours after transfection, live cell imaging was performed on a Zeiss LSM880 confocal equipped with Focus Controller 2, with cells maintained at 37°C and 5% CO_2_ using the TempModule S1 (800-450000, Pecon) and CO_2_ Module S1 (810-450001, Pecon). Images were acquired every 15 minutes for 65 hours using a Plan-Apochromat 20x/0.8 M27 objective at Lancaster University. To determine cell cycle parameters the daughters of a mitosis were tracked in FIJI, either until they completed two full cell cycles or for the duration of the time-lapse.

### RNA extraction and RNA-sequencing library preparation

At least 5×10^5^ cells were lysed using Trizol reagent, and RNA was extracted with the Directzol™ RNA Mini Prep Kit (Zymo Research, cat. no. R2050) according to the manufacturer’s protocol. 1 μg of RNA was used as input for library preparation. rRNA depletion was performed using the NEBNext rRNA Depletion Kit v2 (NEB, cat. no. E7405), followed by cDNA synthesis, end prep, and library amplification using the NEBNext Ultra II Directional RNA Library Prep Kit for 56 Illumina (NEB, cat. no. E7760). The average fragment length was determined using D1000 ScreenTape (Agilent, cat. no. 5067–5582) with D1000 reagents (Agilent, cat. no. 5067–5583), and sample concentration was measured with the Qubit 3.0 Fluorometer (Life Technologies). Next-generation sequencing was performed by Macrogen, Singapore.

### RNA-seq data analysis

Trim Galore (v0.4.2_dev; https://www.bioinformatics.babraham.ac.uk/projects/trim_galore/) was used to trim paired-end raw sequencing reads. The trimmed reads were then mapped to the human hg19 reference genome (hg19/GRCh37) obtained from iGenomes using the RSEM pipeline (v1.1.11) ^32^. Differential expression analyses were performed using DESeq2 ^33^. Differentially expressed genes (DEGs) were identified based on a 1.5-fold change in expression and an adjusted p-value of less than 0.05. Pathway analysis was conducted using the Metascape package (https://metascape.org/gp/index.html#/main/step1) and gene set enrichment analysis (GSEA) software (https://www.gsea-msigdb.org/gsea/index.jsp) using the Molecular Signatures Database (MSigDB). Pathways were considered significantly enriched if the FDR-corrected p-value was ≤0.05. Volcano plots were generated using packages ggplot2 and EnhancedVolcano in R software (v4.0.5).

### CUT&TAG assay

1×10^5^ of cells were resuspended in a wash buffer (20 mM HEPES-KOH pH 7.5, 150 mM NaCl, 0.5 mM spermidine). Concanavalin-A beads were prepared by washing twice with binding buffer (20 mM HEPES-KOH pH 7.5, 10 mM KCl, 10 mM CaCl2, 1 mM MnCl2). Ten microlitres of Concanavalin-A beads (Bangs Lab, cat no. BP531) were added to each reaction and incubated with rotation at room temperature for 15 minutes. The cell-bead complex was resuspended in 50 μL antibody buffer (20 mM HEPES-KOH pH 7.5, 150 mM NaCl, 2 mM EDTA, 0.5 mM spermidine, 0.05% (w/v) digitonin, 0.1% (w/v) BSA). One microgram of anti-Ty1 primary antibody (ThermoFisher Scientific, cat no. MA5-23513) was added to each condition. The reactions were incubated at 4°C overnight with gentle rocking. After removing the supernatant containing the primary antibody using a magnet, secondary antibody was added in dig-wash buffer (20 mM HEPES-KOH pH 7.5, 150 mM NaCl, 0.5 mM spermidine, 0.05% (w/v) digitonin) and incubated at room temperature for 1 hour with rotation. The complex was then washed twice with dig-wash buffer and resuspended in Dig300 wash buffer (20 mM HEPES-KOH pH 7.5, 300 mM NaCl, 0.5 mM spermidine, 0.01% (w/v) digitonin). One microlitre of in-house pA-Tn5 Transposase was added and incubated at room temperature with constant rotation for 1 hour. The complex was washed twice with Dig-300 wash buffer and resuspended in 300 μL Dig-300 wash buffer with 10 mM MgCl2 and incubated at 37°C for 1 hour for tagmentation. The reaction was then quenched by adding 10 μL of 0.5 M EDTA, 3 μL of 10% (w/v) SDS, and 2.5 μL of 20 mg/mL proteinase K and incubated at 50°C for 1 hour. DNA was purified with Serapure beads and eluted in 0.1x TE buffer. Libraries were prepared by tagging each sample with a unique pair of indexes provided in NEBNext® Multiplex Oligos for Illumina® (NEB, cat no. E7311AVIAL) following PCR (Table 2.8). The libraries were purified with Serapure beads. The average fragment length was measured by D1000 ScreenTape (Agilent, cat no. 5067–5582) with D1000 reagents (Agilent, cat no. 5067–5583). Next-generation sequencing was carried out by Macrogen, Singapore.

### CUT&TAG sequencing analysis

Each sample was sequenced with at least 20 million paired-end reads (150x150). Adapter and quality trimming of the reads were performed using Trim Galore (https://github.com/FelixKrueger/TrimGalore) with the parameters --paired --nextera. The reads were aligned to the human genome hg19 using Bowtie2 (version 2.4.5) with the parameters --end-to-end --very-sensitive --no-mixed --no-discordant --phred33 -I 10 -X 700 ^34^. The Bowtie2 index was obtained from the Illumina iGenome website (https://sapac.support.illumina.com/sequencing/sequencing_software/igenome.html). Unmapped reads were removed using samtools view (https://www.htslib.org/doc/samtools.html) with the parameters -bS -F 0x04. Bigwig files as well as metagene plot were generated using deepTools ^35^. Comparisons between bed files (e.g., common overlapping, unique) were performed using bedtools (v2.30.0) ^36^. For peak calling, macs2 was utilized using the parameters: BAMPE --keep-dup all -g hs –q 0.01 and IgG as a control ^37^. Pie chart was generated were generated using Chipseeker ^38^. Motif enrichment analysis was performed using HOMER ^39^ and BAMM ^40^.

### BioID2 pulldowns

SYCP1 full length, variant 1-954 or deletion D918-942 were cloned into BioiD2-MYC or BioID2-HA plasmids. MCF7 cells were transfected using lipofectamine 2000 according to manufacturer’s instructions, and selected for using 400 μg/ml of G418 (geneticin) for two weeks. MCF7 cell lines stably expressing BioID2 constructs were seeded in four 10cm dishes at 3×10^6^ per dish, for each repeat. Biotin was added at 50μM for 24 hours before cells were lysed and sonicated in the lysis buffer (50mM Tris·Cl pH 7.4, 8M urea, 1x protease inhibitor, 1mM dithioreitol) and 20% Triton X-100. Lysates were centrifuged at 4°C, 13,000g for 10 minutes and the supernatant was incubated with 100μl of Streptavidin conjugated beaded agarose (ThermoFisher) overnight on a roller at 4°C. Beads were washed three times in a wash buffer (50 mM Tris·Cl, pH 7.4, 8M urea) and then spun for 2 mins at 1000g. The supernatant was removed and the pellet was resuspended in 30µl of 25mM biotin in 50mM ammonium bicarbonate, heated at 95°C for 5 minutes before centrifugation for 2 mins at 1000g. Without removal of the supernatant, this process was repeated. 80μl of Laemmli buffer with B-ME was added and the solution heated for 5 mins at 95°C. The solution was frozen at -80°C for 30 minutes before a final centrifugation. The resultant supernatant was removed and sent for mass spectrometry analysis. Prepared BioID2 lysates were analysed using tandem mass tag (TMT) mass spectrometry. For each sample, four technical repeats were sent for analysis at the EMBL proteomics core facility Samples were multiplexed in two batches, with two repeats of each experimental condition on each plate.

### Immunoprecipitation

Cells were seeded in 10cm dishes at 1×10^6^ cells per dish for two days. Lysis buffer (1M Tris pH 7.5, 4M NaCl, 10mM Na_3_VO_4_,1% NP-40 alternative, 0.1M PMSF, 1x protease inhibitor, 1M DTT) was added directly to each plate and cells were scraped before transfer to an Eppendorf tube on ice. After a 30 min incubation lysates were spun at 900g for 4 mins at 4°C and the supernatant was taken. Samples were precleared for 4 hours with the addition of 30μl of PGS on rotation at 4°C. 1μg of anti-SYCP1 antibody (Abcam, 15090) was added and incubated with rotation overnight at 4°C. PGS was added at 30μl per tube for 1 hour at 4°C. Beads were pelleted by centrifugation and the supernatant discarded. The beads were then washed with 1ml of wash buffer A (PBS, 0.2% Triton X-100, 350mM NaCl) followed by centrifugation. The supernatant was removed and washed with 1ml of wash buffer B (PBS + 0.2% Triton X-100) and then centrifuged again. The resultant supernatant had 40μl of Laemmli with B-ME added and was boiled at 95°C for 10 mins. The supernatant was ran on an SDS page gel and blotted using antibodies anti-NFX1 (Abcam, ab176733), anti-NFYA (Rockland, 100-401-100), anti-SP1 (Abcam, ab227383), anti-SP5 (Invitrogen, PA5-103505), anti-KLF3 (Abcam, ab154531) or anti-SP2 (Invitrogen, PA5-35984).

### COS7 culture and transfection

COS 7 cells were maintained in DMEM media supplemented with 10% (v/v) fetal calf serum (FCS), and 100 µg/ml PenStrep (Invitrogen) at 37 °C in 5% CO_2_. Transfections were performed using Lipofectamine 3000 reagent (Invitrogen) following the manufacturer’s instructions.

### Slide preparation and imaging

Cells expressing EGFP-SYCP1 were fixed 24h post-transfection in 100% methanol at - 20°C for 20 min. After fixation, cells were washed 3 times for 5 minutes in PBST (1xPBS, 0.1% Triton-X100), stained with Hoechst (10 µg/ml) for 10 min, and washed once more in PBST. Coverslips were mounted on a slide with 10 µl of Vectashield mounting media (Vector).

The slides were imaged on a Nikon TI2 inverted microscope (Nikon), equipped with a Photometrics Prime 95B camera and Nikon Elements 5.1 software. The images were further processed in ImageJ.

### Cloning

The different SYCP1 constructs were cloned into the pEGFP-C3 vector (Clontech) following the in house SLIC (Sequence and Ligation Independent Cloning) reaction protocol and transformed into chemically competent *E. coli* (DH5α). Plasmid DNA was extracted from the bacterial cultures using the GeneJET Plasmid Miniprep Kit (ThermoFisher Scientific) following the manufacturer instructions. All plasmids were sequenced prior to use.

### Chromatin Immunoprecipitation and Quantitative PCR (ChIP-qPCR)

ChIP-qPCR was performed to determine SP1, SP2, and SP5 binding at gene promoters in MCF7 wild-type (WT) and SYCP1 knockout (KO) cell lines. Cells were seeded at 3.5 × 10⁶ in 2x 100 mm dish and cultured for 3–4 days. Crosslinking was performed with 1% formaldehyde for 7 minutes at room temperature, quenched with 0.125 M glycine, and washed with ice-cold PBS. Cells were scraped, pelleted, and either processed immediately or snap-frozen at −80°C.

Chromatin was extracted through sequential lysis buffers and sonicated (30s on, 20 s off, 10 cycles) to yield fragments of 400–600 bp, as confirmed on an agarose gel after RNase A and proteinase K treatment. For each ChIP, 80 µg of chromatin was incubated overnight at 4°C with 2 µg of antibody pre-bound to 40 µL of Dynabeads Protein G (Invitrogen, 10003D). Antibodies against SP1 (Abcam, ab227383), SP2 (Invitrogen, PA5-35984), and SP5 (Invitrogen, PA5-103505) were used, along with a non-immunised, species matched IgG control. Beads were washed five times with ice-cold RIPA buffer and once with TBS. Chromatin was eluted and reverse crosslinked overnight at 65°C, followed by proteinase K treatment and purification using the GeneJet PCR purification kit (Thermo Scientific, K0701).

Candidate target genes were selected using the UCSC Genome Browser ^41^ for SP1, SP2, and SP5, based on regions with strong predicted promoter occupancy. These were then compared with genes identified by SYCP1 CUT&Tag to identify overlapping targets. Overlapping as well as unique targets for each transcription factor were selected for qPCR analysis. Promoter regions 1000 bp upstream of the transcription start site were retrieved for each target gene. Transcription factor binding motifs were obtained from the JASPAR ^42^ database and mapped to promoter regions using FIMO ^43^. Primers were designed to amplify 100–200 bp regions encompassing the predicted binding motifs.

qPCR was performed using PowerUp™ SYBR™ Green Master Mix (Applied Biosystems, A25742) on a standard real-time thermal cycler. Each ChIP condition was analysed in three biologically independent replicates. Data were normalised to input DNA using the percentage input, and fold enrichment was calculated relative to IGG to correct for non-specific binding. Statistical comparisons between WT and KO samples were made using two-tailed unpaired Welches t-test, with significance defined as p < 0.05.

### Statistics and reproducibility

All data was first tested for parametric distribution and analysed with GraphPad PRISM software. Parametric data was analysed using appropriate *t*-tests or ANOVA with Bonferroni’s comparison test for multiple group comparisons. Non-parametric data was analysed using Mann–Whitney or Wilcoxon signed-rank test. By convention, \*\*\**p*-values < 0.001, \*\**p*-values < 0.01, and \**p*-values < 0.05. Number of experiments is indicated for each figure where experiments with internal replicates are shown ±SEM while experiments where that was not possible are shown ±SD.

Survival analysis of TCGA cancer patients was performed on samples of primary tumors. The patients were ordered according to *SYCP1* expression and the cut-off between 1st and 3rd quantile was used to divide patients into *SYCP1* high and low expressing subgroups.

## Author contributions

LBC - generated data in Figs 1e, 2d, 3e, 5b, d-i, 6a, c-d, 7-10, Sup fig 7, took part in study design and manuscript writing

CFJ - generated data in Fig 1c

OG - generated data in Figs 3-4 and Sup fig 5

HZ – generated supp figs 1-2

MP - generated data in Fig 6b, generated reagents used in all plasmid based experiments, took part in study design

ABD – performed computational analysis that provided preliminary data for some of the investigation

SIF - generated data in Figs 1d, 2, 5c and sup fig 4

HM, TA, FAB - generated data in Fig 5a

CG – generated data in Sup Fig 3 MG - generated Fig 1a

ORD – provided biochemical expertise, generously shared data and reagents, took part in regular meetings on study progress and design, participated in manuscript writing and editing, provided creative input to the project

WWT – supervised the RNAseq and Cut&Taq data analysis and proof read the RNAseq and C&T manuscript section

ULM – secured funding, designed the study, led the study, wrote the manuscript,

## Supplementary figures

**Supplementary Figure 1.**
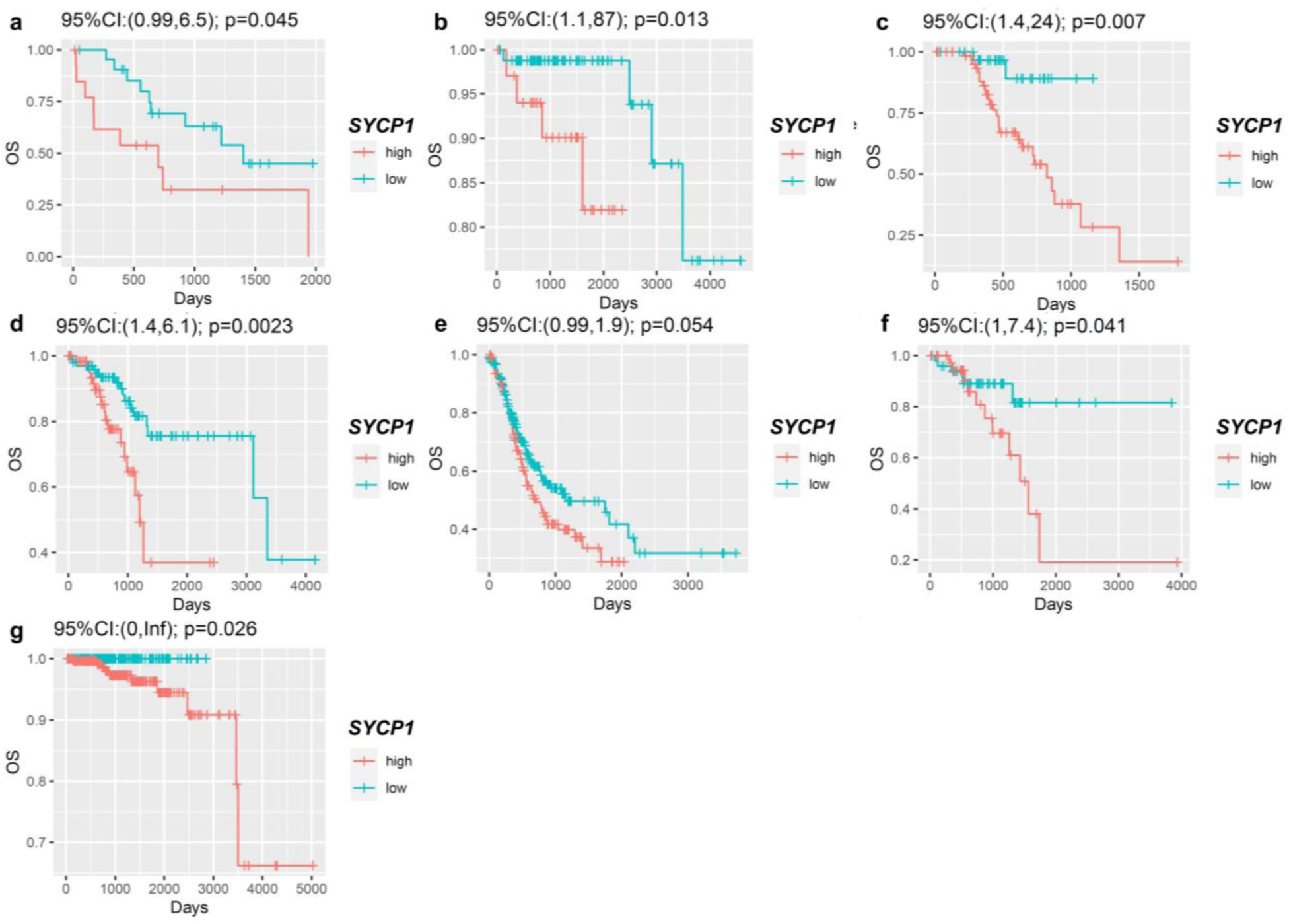
TCGA patients with high SYCP1 expression have shorter overall survival. (**a**) cholangiocarcinoma HR 2.5, (**b**) thymoma HR 9.7, (**c**) skin cutaneous melanoma HR 5.8, (**d**) uterine corpus endometrial carcinoma HR 3, (**e**) stomach adenocarcinoma HR 1.4, (**f**) rectal adenocarcinoma HR2.7. p-value was calculated by log-rank test. HR: hazard ratio, (**g**) prostate adenocarcinoma HR 4.2^e+08^

**Supplementary Figure 2.**
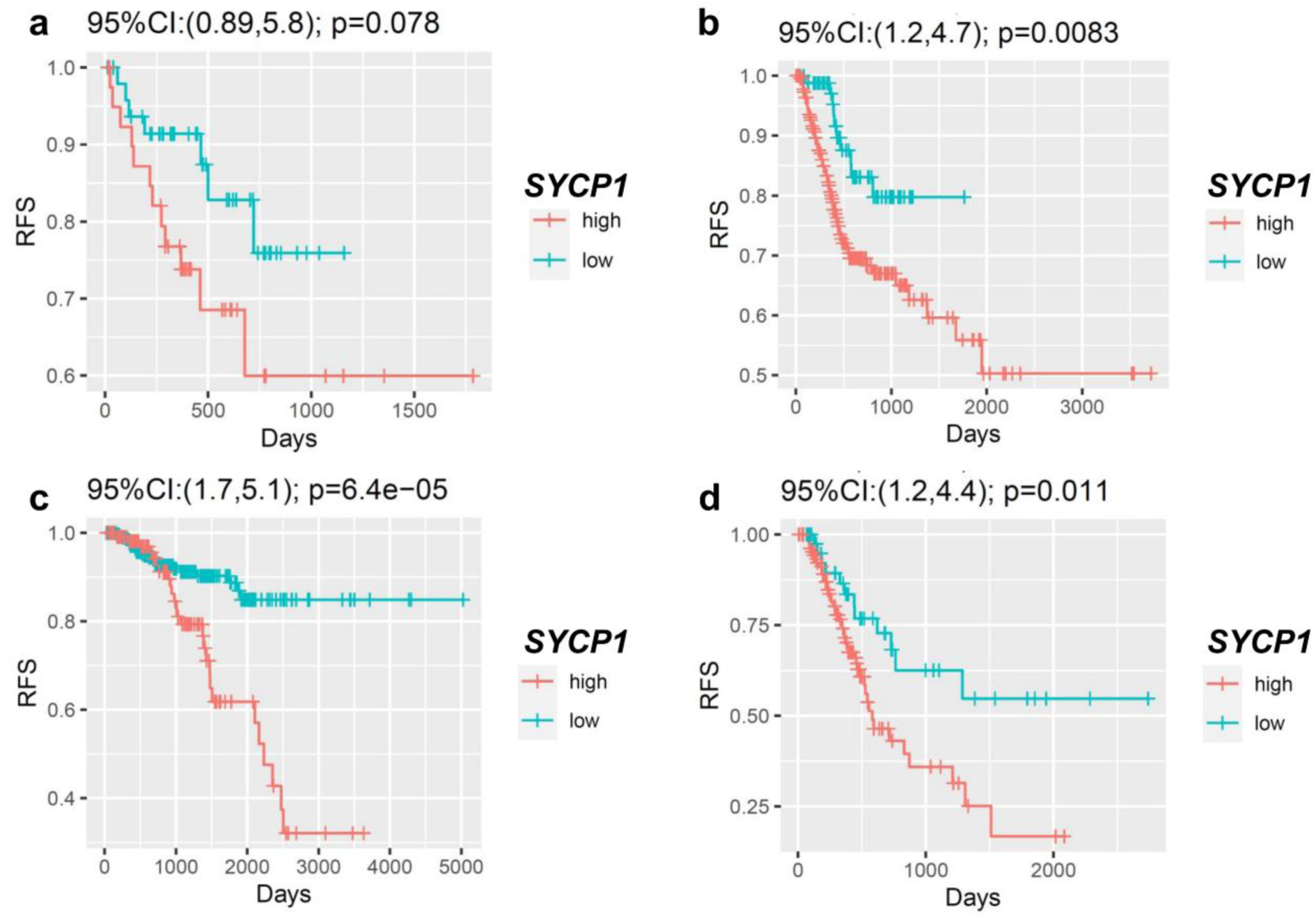
TCGA patients with high SYCP1 expression have shorter relapse free survival. (**a**) skin cutaneous melanoma HR 2.3, (**b**) stomach adenocarcinoma HR 2.4, (**c**) prostate adenocarcinoma HR 2.9, (**d**) pancreatic adenocarcinoma HR 2.3. p-value was calculated by log-rank test. HR: hazard ratio.

**Supplementary Figure 3.**
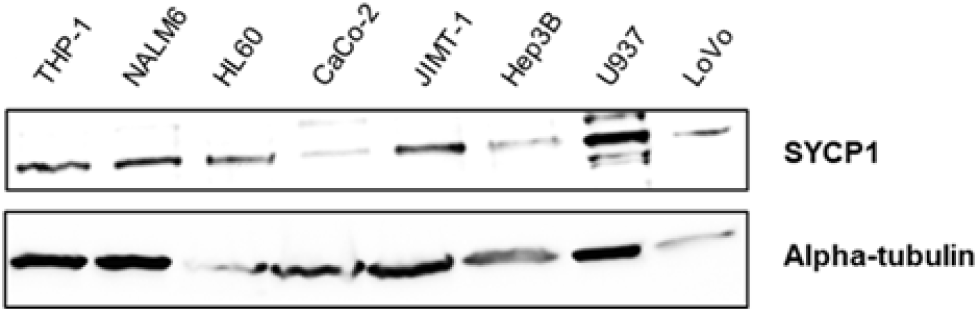
SYCP1 is commonly expressed in cancer cell lines. SYCP1 protein levels across cancer cell lines.

**Supplementary figure 4.**
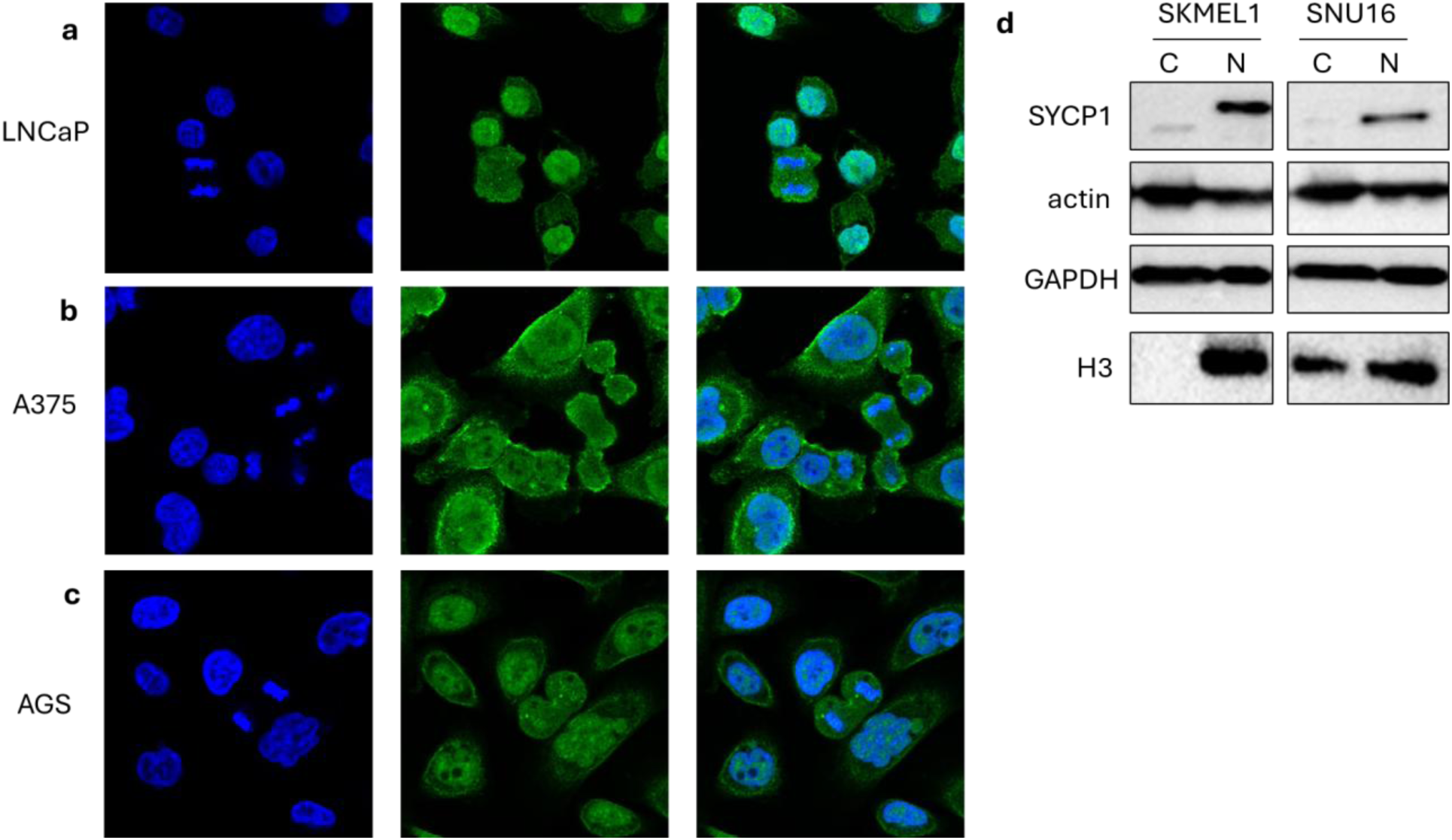
SYCP1 is a nuclear protein in cancer. (**a**) prostate cancer LNCaP cells, (**b**) stomach cancer A375 cells, (**c**) stomach cancer A375 cells. SYCP1 immunolabelled in green with DNA DAPI stained in blue, (**d**) cell fractionation of SKMEL1 melanoma cells and SNU16 stomach cancer cells.

**Supplementary figure 5.**
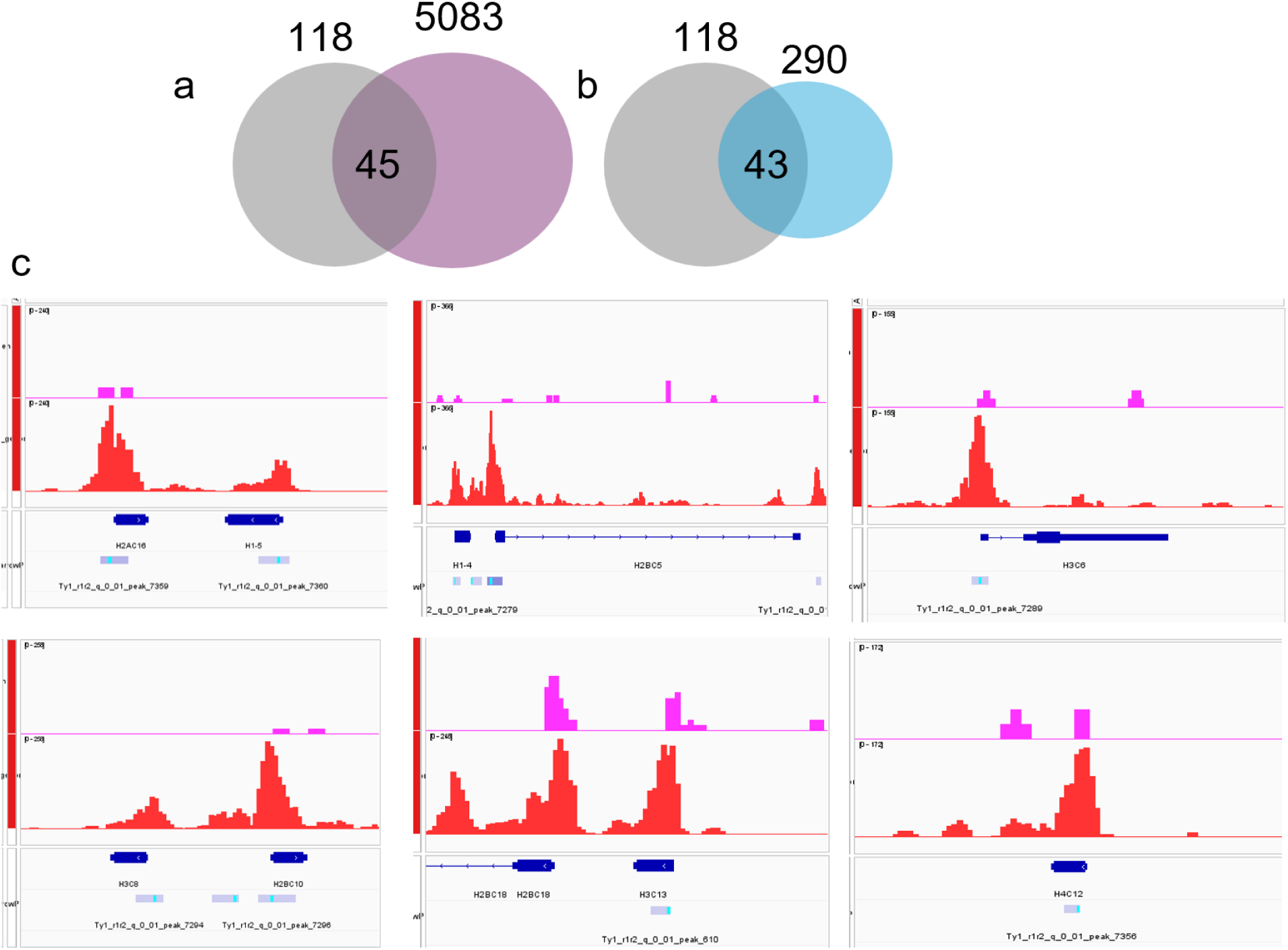
SYCP1 preferentially binds to promoters and regulates expression of histone genes. (**a**) comparative analysis of 5083 genes with SYCP1 promoter binding vs histone genes using all histone genes from HUGO, Parameters: 45, 5083, 118, 20000; expected number of successes = 29.9; results are over enriched 1.5 fold compared to expectations; hypergeometric p-value = 0.0015. (**b**) comparative analysis of 290 genes with SYCP1 promoter binding that were also significantly downregulated upon SYCP1 knock-out vs histone genes using all histone genes from HUGO, Parameters: 45, 290, 118, 20000; expected number of successes = 1.71; results are over enriched 26.3 fold compared to expectations; hypergeometric p-value = 1.8e-52. (**c**) SYCP1 histone peaks.

**Supplementary figure 6.**
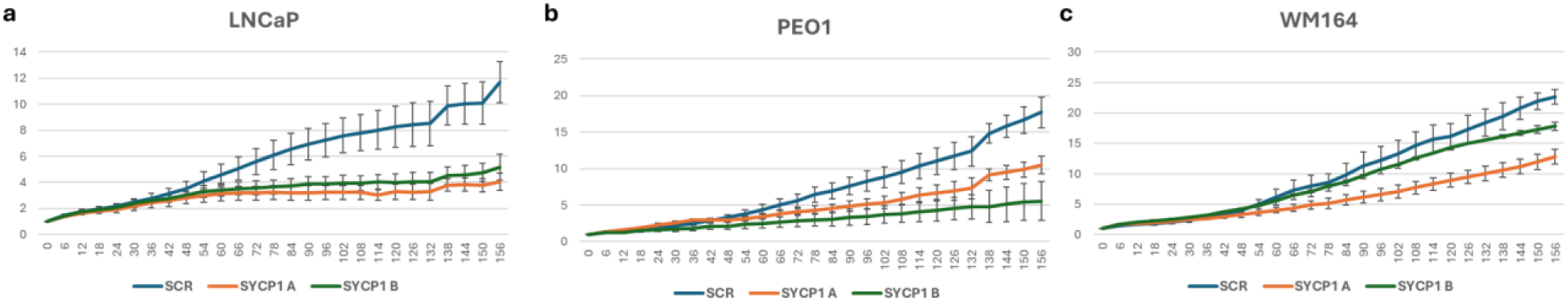
**SYCP1 silencing abrogates cancer cell growth**. (**a-c**) Cells were seeded with non-targeting siRNA (SCR) and two SYCP1 targeting sequences (SYCP1 A and B). Cell growth was monitored using incucyte for 156hrs and is presented as fold change over control SCR treated cells. (**a**) LNCaP prostate cancer cells, (**b**) PEO1 ovarian cancer cells, (**c**) WM164 melanoma cells.

**Supplementary Figure 7.**
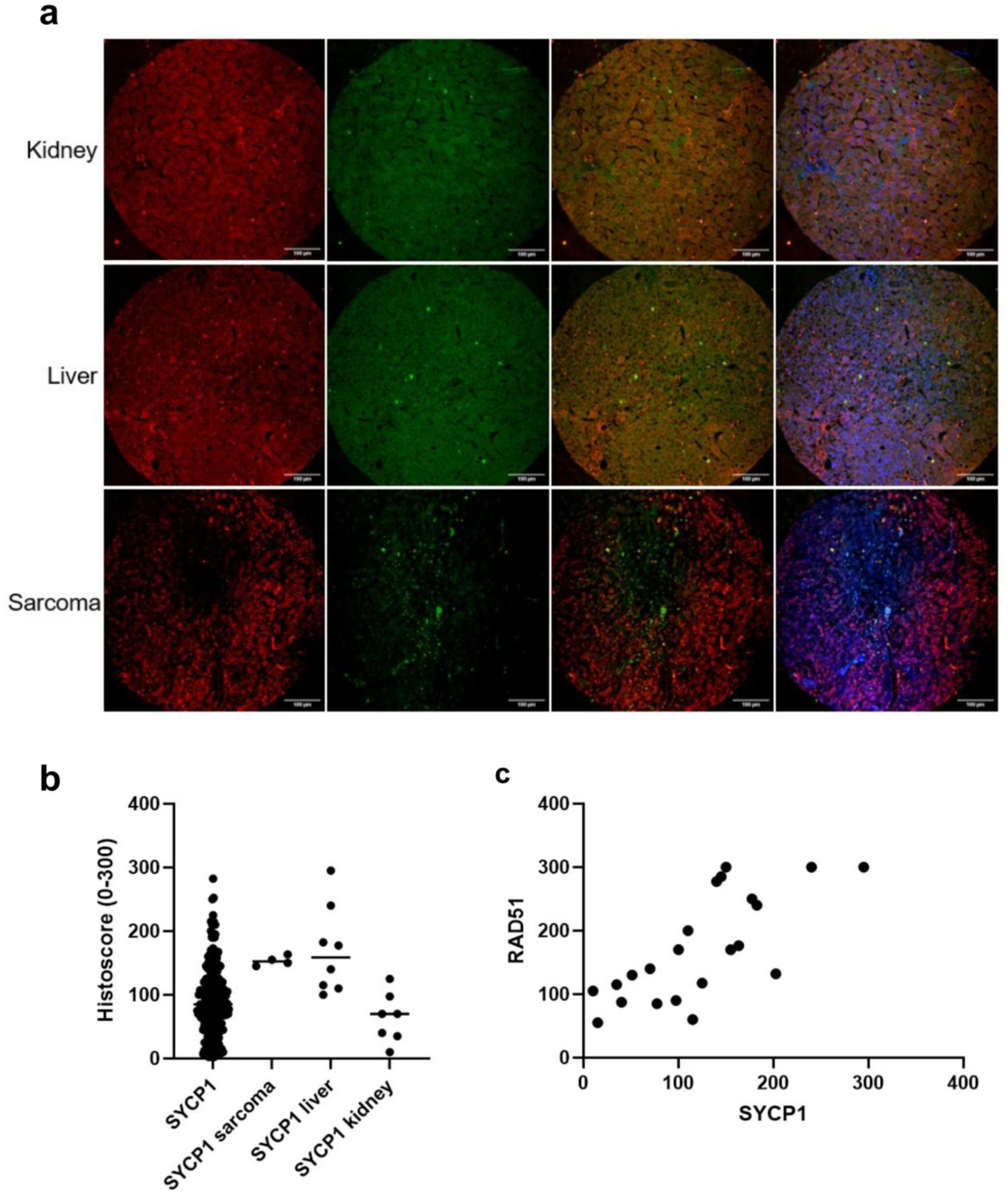
SYCP1 colocalises with RAD51 in kidney, liver and sarcoma patient. (**a**) TMAs from kidney, liver and sarcoma cancer patients were stained for both RAD51(green) and SYCP1 (red) (**b**) SYCP1 histoscore distribution by cancer type.(**c**) RAD51 and SYCP1 have a positive correlation in the cohort (p<0.0001).

## Acknowledgements

Authors would like to thank Dr Richard Mort (Lancaster University) for generous gift of MCF7 Fucci cell and Dr Gerben Vader (PMC, Utrecht) for insightful discussion. ULM and LB were supported by MRC (MR/X00855X/1), ULM is also funded by North West Cancer Research (RDG2021.15) and the MRC with CF (DiMeN DTP2), as well as by a European Molecular Biology Organization Young Investigator Network Grant and Sonata Bis. W-W.Tee is supported by the National Medical Research Council [NMRC/OFIRG21nov-0027 and OFIRG24jul-0094]; A*STAR Biomedical Research Council, Central Research Fund, Use-Inspired Basic Research (CRF UIBR) and Competitive Research Programme (CRP) [NRF-CRP31-0024]. O.R.D. is a Wellcome Senior Research Fellow (Grant Number 219413/Z/19/Z) and was supported by the Light Microscopy Core of the Wellcome Discovery Research Platform for Hidden Cell Biology (226791). Authors would like to thank the EMBL Proteomics facility for exceptional support, all MS/MS analysis were performed at EMBL supported by ULM EMBO Young Investigator Network IG grant.

